# On the genetic consequences of habitat contraction: edge effects and habitat loss

**DOI:** 10.1101/2022.10.25.513679

**Authors:** Gabriele Maria Sgarlata, Tiago Maié, Tiago de Zoeten, Rita Rasteiro, Lounès Chikhi

## Abstract

Natural climate change and recent anthropogenic activities have largely contributed to habitat loss and fragmentation across the world, leading to 70% of worldwide remaining forests to be within 1 km of forest’s edges (Haddad et al., 2015). Ecological studies have shown that edge-effect influences ecological communities, species richness and abundance across many taxa, contributing to worldwide decline in biodiversity. Since edge-effect reduces species abundance and connectivity, it is also expected to negatively influence species genetic variation. In fact, previous theoretical studies had showed that populations closer to the edges of a finite stepping-stone model tends to have shorter coalescence times, and therefore, lower genetic diversity, than central populations. However, predicting the impact of edge effect on local genetic diversity remains challenging in realistic and more complex habitat fragments, where the additive effect of multiple edges is expected to take place. In the present study we explore the genetic consequence of habitat loss at the scale of a habitat fragment (*patch-scale*), looking at the interplay between *patch-size* and *edge-effect* on spatial genetic diversity. We propose a statistical approach to estimate ‘edge-impacted effective population size’ from habitat cover information and use this measure to predict spatial genetic diversity in both equilibrium and non-equilibrium populations. We address these questions using spatially-explicit simulations and propose a spatially-explicit analytical framework able to model spatio-temporal changes in genetic diversity due to edge-effect and habitat loss.

## Introduction

Throughout natural history, landscapes have undergone through changes in configuration, abiotic characteristics and level of connectivity (Williams, 2003; Hewitt, 2000; Weigelt et al., 2016). Those changes have affected the distribution of species and ecosystems worldwide, at different rate and extent (Graham et al. 1996; Williams et al., 1998; Lyons 2003, 2005). More recently, human activities had dramatically modified and destroyed natural environments (Barlow et al., 2016; Song et al., 2018), with the consequent decline in global biodiversity (Pimm et al., 2014; Newbold et al., 2015; Ceballos et al., 2015, 2017). Habitat loss and fragmentation (HL&F) has been recognized as one of the major threats for biodiversity (Groombridge, 1992; Pereira et al., 2010; Tittensor et al., 2014). Haddad et al., (2015) estimated that world remnant forests are greatly affected by tree cover loss and isolation, leading to about 70% of worldwide forests being at 1 km from an edge. Vancutsem et al., (2020) have shown that 17% of the tropical moist forests have disappeared between 1990 and 2019, and predictive models suggest that by 2050 undisturbed forests will disappear entirely in large tropical humid regions.

HL&F is a process characterized by loss of habitat amount and the breaking apart of habitat in smaller patches (Curtis 1956; Moore 1962; Fahrig, 2019). The responses of biodiversity to HL&F are different at the scale of the *landscape* (multiple habitat fragments) and at the scale of the *patch* (habitat fragment) (Pardini et al., 2010; Didham et al., 2012; Fletcher et al., 2018; Fahrig, 2019). At the *landscape*-scale there are two processes that may occur: *ĩ*) removal of habitat (habitat loss); and *ii*) an increase in the number of habitat patches within a landscape (habitat fragmentation *per se*). At this scale the variables that are quantified for describing the mechanisms underlying a biological response (ecological or genetic) to HL&F are ‘habitat amount’ and ‘habitat configuration’ (a measure of fragmentation; e.g., number of patches). At the *patch*-scale only habitat removal can occur, since by definition a habitat patch cannot be described as ‘fragmented’ (broken apart), as it represents the smallest unit of study in a fragmented landscape (Fahrig 2003; Fahrig 2017). Therefore, the variables that matter at the *patch*-scale are size (spatial extent of continuous habitat), isolation (distance from the nearest patch) and edge-effect (changes in habitat quality in proximity to the boundaries between habitat and non-habitat). More generally, this means that *patch*-scale and *landscape*-scale mechanisms are not necessarily interchangeable, although related, and they may need first to be investigated separately (Ruffell et al., 2016; Fletcher et al., 2018).

Although the consequences of habitat loss and fragmentation on biodiversity have been largely investigated over the last decades (see references in Fletcher et al., 2018), some controversies still exist. There is a general agreement that habitat loss has detrimental effect on biodiversity (Brooks et al., 2002; Schipper et al., 2008; Fletcher et al., 2018; Fahrig, 2017). In contrast, the extent to which habitat fragmentation has negative effect on biodiversity is still debated (Fahrig et al., 2019; Fletcher et al., 2018). For instance, a recent review of the literature by Fahrig et al., (2017) found that ecological responses to habitat fragmentation *per se* (fragmentation independent of habitat amount) were in >70% of study cases non-significant, and, when significant usually positive. In contrast, a later study by Fletcher et al., (2018) has argued that data from literature rather support a significant negative effect of habitat fragmentation *per se* on biodiversity. This controversy revealed several limitations in HL&F research that need to be overcome in order to resolve the scientific controversy and improve our understanding of biodiversity (Fletcher et al., 2018). We mention some of these limitations listed in Fletcher et al., (2018). First, habitat loss and habitat fragmentation are often correlated processes, that is, one rarely occurs without the other. Statistical analyses that do not explicitly take into account the collinearity of such processes can draw misleading conclusions on the effect of HL&F on species ecology (Didham et al., 2012). Second, ecological responses to HL&F are difficult to generalize across taxa and ecosystems. For instance, a positive response to HL&F by a given species can actually go in favor of a detrimental effect of HL&F, if this species is generalist or invasive (Pfeifer et al., 2017; Fletcher et al., 2018). Third, habitat and non-habitat categorization need to be carried out according to the species ‘perspective (Betts et al., 2014). Ultimately, HL&F processes have to be regarded as spatio-temporal processes, where different mechanisms may act depending on the spatial scale, generating ecological responses that rely on complex temporal dynamics (Hylander and Ehrlé, 2013; Ryo et al., 2019). Therefore, reconciling the scientific disagreement requires research that goes beyond statistical and correlative approaches, and focuses more accurately on the mechanisms acting across spatial-temporal scales (Fletcher et al., 2018).

At the *landscape*-scale, Fahrig, (2017) meta-analysis suggests that habitat fragmentation *per se* can have either non-significant (>70%), positive (>20%) or negative (<10%) effect on biodiversity (species abundance, occurrence and richness). Fahrig L. concluded that perhaps, habitat fragmentation *per se* does not influence biodiversity as much as loss of habitat amount. Accordingly, data on 5675 species from eight taxonomic groups (plants, fungi, gastropods, insects, amphibians, reptiles, birds and mammals) suggest that ‘habitat amount’ has a stronger negative effect on ecological response (species density; number of species in a plot of fixed size) than measures of habitat fragmentation *per se* (patch size and isolation) (Watling et al., 2020). These results have been interpreted as supporting the ‘Habitat Amount Hypothesis’ (HAH), which states that species density increases proportionally to the amount of habitat present in the area where individuals of a given species are likely to arrive (Fahrig 2013; Martin and Fahrig, 2012). This area can extend beyond the boundaries of the habitat patch from which ecological data have been retrieved, thus suggesting that patch-size may not be a full predictor of the measured ecological response. The HAH is still debated, in part because some empirical studies have demonstrated the importance of mechanisms occurring at the *patch*-scale, such as edge-effects and patch-size, to explain ecological patterns (e.g., Hanski et al., 1995; Ries et al. 2004; Prevedello and Vieira, 2010; Hanski et al., 2013; Rybicki and Hanski, 2013; Haddad et al., 2015; Pfeifer et al., 2017; Ries et al., 2017; Haddad et al., 2017; Betts et al., 2019; Bueno and Peres 2019). On the last point, indeed Watling et al., 2020 states that HAH should be seen as a parsimonious approach, where *patch*-scale variables (patch area and isolation) are replaced “with a single measure of habitat amount” that provide “a useful approximation for describing the key landscape driver of species density” in the study area. Therefore, if HAH represents a “useful approximation” of *patch*-scale processes occurring in a fragmented landscape, understanding processes at the *patch*-scale would i) provide accurate insights on the HL&F mechanisms impacting biodiversity; ii) clarify possible controversies; and ii) elucidate how and to which extent extrapolate *patch*-scale mechanisms at a larger spatial scale (*landscape*-scale). In the present study we investigate two of the three main mechanisms occurring at the *patch*-scale: *patch-size* (habitat amount) and *edge-effect*. The meta-analysis of Watling et al., (2020) has shown that *patch-size* does significantly explain species density response (mean effect size = 0.351 ± 0.007), although at a lower extent than *habitat amount* computed at the *landscape*-scale (mean effect size = 0.606 ± 0.007). Previous works have shown that also *edge-effect* has great influence on ecological communities (Ries et al. 2004; Fletcher, 2005; Pfeifer et al., 2017; Ries et al., 2017). For instance, Pfeifer et al., (2017) have developed a statistical approach that quantify additive multiple edge-effect from landcover data (‘Edge Influence’) and proposed a simple measure (‘Edge Sensitivity’) estimating the sensitivity of a given species to edges. Using these statistics, Pfeifer et al., (2017) assessed edge-determined changes in abundance among 1,673 vertebrate species, showing that 85% are affected either positively (46%) or negatively (39%) by edge-effect. The species negatively impacted by edge-effect have 3.7 times more probability to be listed as threatened (according to IUCN classification), whereas species with positive response are typically invasive or generalist species. These studies provided clear evidence on the importance of *patch*-scale processes on ecological response.

Species ecology has a direct impact on species genetic diversity and differentiation. Effective population density (*D_e_*) and dispersal (σ_e_) are two variables typically included in population genetic models, able to capture some of the direct consequences of species ecology on neutral genetics (reviewed in Cayuela et al., 2018). D_e_ and σ_e_ are often different from census population density (D) or dispersal (σ) (e.g., Slatkin, 1987; Broquet and Petit, 2009; Hellegren and Galtier, 2016). This is because D_e_ and σ_e_ are proxies for processes involved on the reproduction of individuals and inheritance of genes, whereas D and σ are estimates based on the actual observation of individuals, which may not all contribute equally to the genetic pool of the population (e.g., Cayuela et al., 2018). Beyond species ecology, genetic patterns are also influenced by the molecular properties of DNA, such as mutation rate (*μ*) and recombination rate (*r*). The commonalities between genetic and ecological processes suggest that HL&F may affect genetic diversity similarly to how it affects species ecological response (e.g., abundance). On the other hand, discrepancies between ecological and genetic processes (e.g., mutation rate) could point out to possible differences on the effect of HL&F on ecological and genetic responses.

Several empirical and theoretical studies have investigated the consequences of HL&F on population genetic variation. Meta-analyses on *landscape*-scale studies around the world and across several taxonomic groups have shown consistent evidence of an overall negative effect of anthropogenic disturbance and HL&F on species genetic diversity (Lino et al., 2019; Almeida-Rocha et al., 2020; Monteiro et al., 2019; Miraldo et al., 2016; Schlaepfer et al., 2018; Vranckx et al., 2012; Rivera-Ortíz et al., 2015; Aguilar et al., 2008; Honnay & Jacquemyn, 2007), although some have observed either neutral or non-significant effect (land-use intensity and human density; Milette et al., 2020). The recent global analysis of Almeida-Rocha et al., (2020) investigated the effect of *landscape*-scale and *patch*-scale disturbance on species genetic responses. They stated that *landscape*-scale measures of habitat loss and reduction in connectivity were the factors with most detrimental effect on genetic diversity (respectively, forest cover effect size: −0.67 [−0.89, −0.44]; fragments isolation effect size: −0.65 [−1.28, −0.02]). The *patch*-scale processes with a significant negative effect on genetic diversity were ‘local quality’ (effect size: −0.08 [−0.38, −0.21]) and ‘patch size’ (effect size: −0.42 [−0.65, −0.20]), although, we could consider also ‘connectivity’ as a *patch*-scale process, since Almeida-Rocha et al., (2020) defined it as the level of physical isolation between habitat fragments (see definition of *patch-isolation* in Fahrig, 2019). It is worth to note that in Almeida-Rocha et al., (2020), *edge-effect* did not appear to have a significant influence on species genetic response. This is probably due to the small sample size of *edge-effect* genetic studies included in the meta-analysis (N = 1; Quevedo et al., 2013), which is probably related to the fact that no much research has been dedicated to such process yet (see Radespiel et al., 2019 for an exception). This reflects a general tendency in landscape genetic literature, where most studies are focused on the effect of HL&F on population genetic differentiation at the *landscape*-scale, and very few on population genetic diversity at the *patch*-scale (Storfer et al., 2010; DiLeo and Wagner, 2016).

Theoretical and simulation-based studies on the genetic consequences of HL&F are also skewed towards patterns of population genetic differentiation at the *landscape*-scale (Cushman et al. 2012; Mona et al., 2014; Jackson and Fahrig, 2014; Jackson and Fahrig, 2016). However, there are some exceptions. The earlier theoretical studies investigating the impact of *edge-effect* on genetic diversity date back to the work of Maruyama, (1972) and Tajima, (1990), who performed numerical computations of the expected amount of genetic diversity in several structured models. They both showed that lower dispersal rate generates lower genetic diversity, and assuming that edge populations experience lower dispersal rate than central populations, then it is expected that edge populations would have lower genetic diversity (Tajima, 1990). Later theoretical works have confirmed Tajima’s predictions, showing that populations closer to the edges of a finite habitat (stepping-stone model) tends to have shorter coalescence times, and therefore, lower genetic diversity (Hey, 1991; Herbots, 1994; Wilkins and Wakeley, 2002; Wilkins, 2004). In particular, Wilkins (2004) has derived explicit expressions of the distribution of coalescence times for edge-impacted populations in a rectangular habitat. Following theoretical studies have also pointed out the impact of habitat boundaries on genetic variation (e.g., Leblois et al., 2006; Duforet-Frebourg and Slatkin, 2016). Leblois et al., (2006) performed spatially-explicit simulations to study the effect of habitat contraction on population genetic diversity. They observed that samples collected across the whole habitat were showing lower observed heterozygosity than samples collected exclusively from the centre. They imputed these results to the effect of edges at the boundaries of the simulated scenario. Recently, Ringbauer et al., (2017) have proposed a framework that model the effect of a linear barrier on the distribution of coalescence times, on the probability of identity-by-descent, and therefore on genetic diversity. The barrier would be equivalent to the edges of a habitat patch. However, similarly to Wilkins, (2004), the use of such model might become computationally expensive in realistic and more complex habitat fragments, where the additive effect of multiple edges (= multiple barriers) is expected to take place (Pfeifer et al., 2017).

In the present study we explore the genetic consequence of habitat loss at the *patch*-scale, looking at the interplay between *patch-size* and *edge-effect* on spatial genetic diversity. We propose a statistical approach to estimate ‘edge-impacted population size’ from habitat cover information (proxy for De), and use this measure to predict spatial genetic diversity in both equilibrium and non-equilibrium populations. Our approach was in part inspired by the ecological measure of ‘Edge Influence’ introduced by Pfeifer et al., (2017), and by previous works on landscape genetics that uses ‘neighbourhood-level’ (or ‘node-based’) analysis to correlate habitat amount with genetic diversity (e.g., Keyghobadi et al., 2005). Pfeifer et al., (2017) results show that, across many taxa, ‘Edge-Influence’ statistics correlates with species abundance (ecological response), but remain currently unknown the performance of this approach on predictions of edge-determined changes in genetic diversity (genetic response). In conclusion, this work is set within the context of a true integration of population genetic and landscape ecology methods for landscape genetics, since often these disciplines use similar concept but little theoretical overlap (Storfer et al., 2006; Wagner and Fortin, 2013).

## Material and Methods

### Background

Pfeifer et al., (2017) have developed a metric called ‘Edge Influence’ (*I*) for quantifying the cumulative effects of multiple edges, including edge shape and patch-size, from maps of habitat cover (e.g., percentage tree cover). This measure provides quantitative information of *I* for a focal point in the map, and thus allow to relate ‘edge-influence’ with some other variable of interest, such as species abundance. It is defined as:

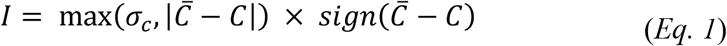

where *C* is the habitat cover at the focal point and, 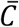 and *σ_c_* are, respectively, the average and standard deviation of habitat cover within an area *A* around the focal point. In Pfeifer et al., (2017), 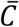 and *σ_c_* are estimated using a Gaussian filter, that is by weighting the contribution on 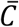 and *σ_c_* of each data point within *A* by their distance from the focal point. The Gaussian filter weights each data point according to a Gaussian distribution:

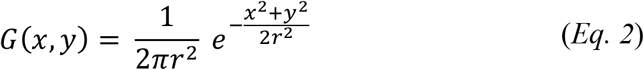

where *x* and *y* are the distances, respectively, on the *x* and *y* direction from the focal point, and r the radius of the area *A*. Note that the Gaussian weight depends only on geographical distance and radius *r*. The value of I corresponds to the maximum between the regional (*σ_c_*) and the point heterogeneity 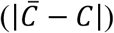, with a sign given by 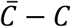, which is negative if the focal point is near the matrix (non-habitat) and tends to zero while moving away from the matrix, toward the interior of the habitat.

### Edge-corrected Effective Population Size

Similar to Pfeifer et al., (2017), we propose an approach that can quantify the additive effect of multiple edges at each point in space within a 2D stepping stone model with irregular shape. Such analysis provides an ‘edge-influence index’ for each deme in the grid, that is used to correct local carrying capacity and obtain an estimator of ‘edge-corrected population size’ (*N_edge_*). The ‘edge-influence index’ can be interpreted as a measure of connectivity within habitat patch, assuming that proximity to habitat edges determines the extent to which each deme can receive alleles from any direction in the 2D space. Deme ‘edge-influence index’ was calculated from a grid representing the shape and the habitat structure (carrying capacity) of the patch. Carrying capacity information within habitat patch are considered to be a proxy for local census population size of the studied species, and could be retrieved, for instance, from data on habitat cover (e.g., percentage of tree cover, NDVI). For each deme, we computed the weighted mean of deme population size from the total number of demes included within a square matrix with side length ‘*R*’ centred in the focal deme. The weights of the mean were defined using the correlation coefficient of allele frequencies between pairs of demes at a given distance ‘*k*’ as detailed in Kimura and Weiss, (1964):

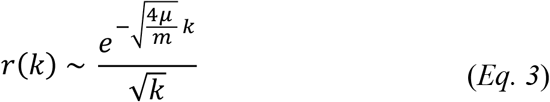

where *μ* is the per site mutation rate and *m* is the deme dispersal rate.

Thus the ‘edge-influence index’ of a deme with spatial coordinates ‘*x, y*’ (*E_x, y_*) is computed as:

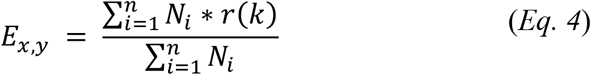

where *N_i_* is the population size of deme *i* and *n* is the total number of demes that fall within the square matrix area with side length ‘*R*’, centred in ‘*x, y*’. The smallest *R* that maximises the fitting between genetic diversity and ‘edge-corrected population size’ is selected as being the optimal radius, since it limits computational cost while being sufficiently accurate (see *Results*). The ‘edge-influence index’ for each deme is then normalized with respect to the ‘edge-influence index’ value in a 2D homogenous habitat (equal *N_i_* across demes) with no boundaries 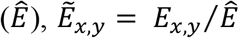. We then obtain the ‘edge-corrected population size’ of each deme in the habitat by 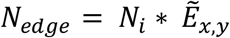. A graphical representation of this approach is presented in Supplementary Appendix S1: Figure S1.

### Two-dimensional spatial model at equilibrium

Similar to Barton et al., (2013), we model the observed heterozygosity (*H*_0_) within a sub-population (deme) in a 2D stepping-stone by using the probability of identity-by-descent of two alleles randomly sampled within the same sub-population (*F*_0_):

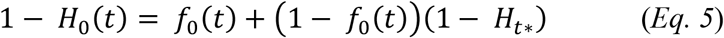

where *H*_0_(*t*) is the observed heterozygosity at time *t, f*_0_(*t*) is the probability of identity-by-descent at time *t* and *H_t*_* is the heterozygosity at a given time *t^*^*, a timepoint beyond which geography (sampling location) is assumed to have little or no impact on the distribution of lineages. In other words, *Eq. 5* states that two alleles randomly sampled within subpopulations can be homozygous either due to identity-by-descent (first term) or by chance (second term). Given a sampling area *A* which diameter is much lower than 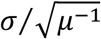, where *σ* is the dispersal distance of a studied species and *μ* is the per site mutation rate, Barton et al., (2013) define *t^*^* as a time satisfying Diameter(A)^2^/σ^2^ ≪ *t^*^* ≪ *μ*^−1^. Assuming a low mutation rate, in the time window between the present and *t^*^* in the past, no new mutations have appeared and a considerable proportion of alleles are very closely related (‘fast local coalescence’) (Barton et al., 2013). As a consequence, the genetic diversity observed at this spatial and temporal scale refers to mutations that occurred more than *t^*^* generations ago. Similarly, Wilkins (2004) defines a ‘transition time’ *τ* between two ‘phases’ of the coalescence process in spatially structured populations: ‘*scattering-phase”* (dependent on sampling location) and ‘*collecting-phase”* (independent from sampling location), respectively. Wilkins (2004) results were based on previous studies showing that the genealogical process in structured population can be modelled using the ‘separation-of-timescales’ approach (Wakeley, 1999; Wilkins and Wakeley, 2002). According to this approach, the genealogy of genes in a structured model can be divided in two time ‘timescales’: ‘ *scattering phase*’ and ‘ *collecting phase*’. The ‘ *scattering phase*’ corresponds to the distribution of coalescence times in the recent past, which is dependent on the original sampling location and on the details of the structured model (dispersal rate, density, drift). In a spatially structured model, it can be modelled as a random walk of lineages, which coalesce once they are in the same deme, at a rate given by the inverse of the deme population size. The ‘ *collecting phase*’ corresponds to the distribution of coalescence times in the ancient past, which is independent from the original location of the sampled genes. In this phase, coalescence rate is dependent on the total population size of the 2D stepping-stone, given that it mostly includes lineages that have crossed the population multiple times since the time to their most common ancestor (Barton and Wilson, 1995). Referring to *Eq. 5*, the ‘*scattering phase*’ is modelled by the probability of identity-by-descent (first term), whereas the ‘*collecting phase*’ is represented by the heterozygosity at time *t^*^* (second term).

At mutation-drift equilibrium (*t* → ∞), the probability of identity-by-descent of two alleles randomly sampled within the same deme, can be written using the Wright-Malécot approximation (Wright 1943; Malécot 1948; Barton et al., 2002):

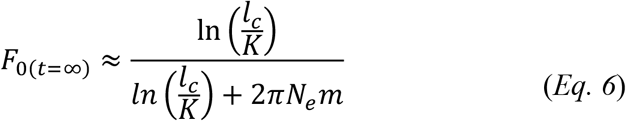

where 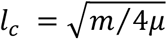 represents the balance between mutation and dispersal, *m* is the deme dispersal rate in a 2D stepping stone model and *K* is the ‘local scale’ constant, which in the case of a 2D stepping stone model is equal to 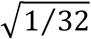. If we assume a dispersal distribution of Gaussian form (as it might be the case to assume with real data), *m* can be replaced by 2*σ*^2^ and *K* = *σ*.

As mentioned above, the heterozygosity at time *t^*^* is determined by lineages coalescing more than *t^*^* generations ago (‘*collecting-phase*’) and that had enough time to cross the whole population (Barton and Wilson, 1995). It has been shown that, under a range of parameter values, the genealogy of most lineages in the ‘*collecting-phase*’ can be approximated by the coalescence process in a panmictic population, which size is equal to the total population size in a 2D stepping stone model (Cox and Durrett, 2002; Wilkins, 2004). This measure relates also to the average coalescence time of two alleles randomly sampled within the same deme (Strobeck 1987; Slatkin, 1991; Hey, 1991). Therefore, we can use the expression for the expected coalescence time of a panmictic population of size equal to the whole population to derive *H_t*_*. Since in the present study we are measuring heterozygosity using microsatellite data, *H_t*_* is written as the expected heterozygosity at mutation-drift equilibrium under a Stepwise Mutation Model (microsatellite) (Kimura and Ohta, 1973; Ohta and Kimura, 1973):

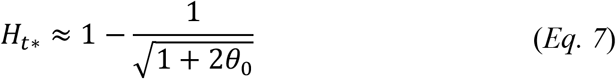

where 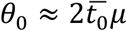 is the expected number of differences between sampled alleles (or population mutation rate) under the infinite site model of mutation and 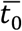 is the expected coalescent times of two alleles sampled from a panmictic population with size equal to the 2D stepping stone model. The expected coalescence time 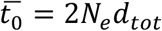 can be written as 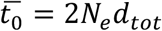. where *N_e_* is the deme effective population size and *d_tot_* is the total number of demes in the population.

So far, these expressions assume homogeneous deme effective population size across the 2D stepping-stone. To model the consequences of edge-effect on deme heterozygosity across a habitat fragment of irregular shape, we compute the edge-corrected probability of identity-by-descent at mutation-drift equilibrium for each deme in the grid by using in *Eq. 6* the estimated *N_edge_* defined in the section above. Then, by rearranging *Eq. 5*, and using *Eq. 6* and *Eq. 7*, we can write the edge-corrected observed heterozygosity as:

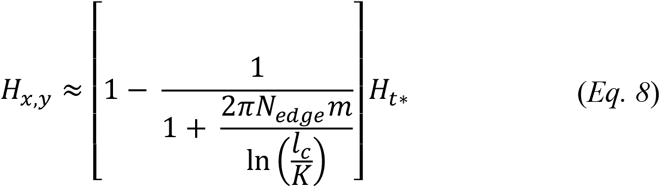

### A statistical approach for testing molecular edge-effect

As an alternative approach to the spatial model presented above, we used a generalized additive model (GAM) to quantify the impact of the ‘edge-corrected population size’ on the observed heterozygosity. The GAM model was chosen because of its flexibility for both linear and non-linear relationships between predictor and response variable, and its capacity to capture patterns that classical linear models would miss (Hastie and Tibshirani, 1987). In particular, it estimates an unknown ‘smooth’ function of some predictor variable using non-parametric approaches to maximize the fitting of the data. We used the *gam* function from the *mgcv* R package (Wood, 2011), setting REML as estimation method for the smooth parameter. We quantified the predictive power of the ‘edge-corrected population size’ by i) measuring the total ‘Deviance explained by the model’, which represents the proportion of the total differences between observed and predicted values; or ii) by quantifying the Normalized Root-Mean-Square Error (NRMSE) between predicted and observed values. NRMSE measures the average squared difference between estimated and observed values, normalized by the range of the observed data: 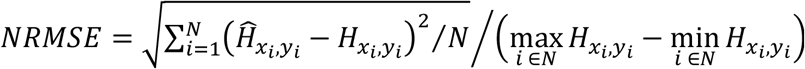. We computed NRMSE for both spatial and GAM models.

### Two-dimensional spatial model at non-equilibrium

We use coalescent theory in a 2D stepping-stone model to describe the temporal dynamic of deme heterozygosity in habitat patches undergoing contraction. Our model considers a 2D stepping-stone model that has been evolving for *τ*_2_ generations. If we consider the present time as *t* = 0, the individuals forming the initial population *τ*_2_ generations ago are randomly sampled from a panmictic model with size *N*_0_ and heterozygosity given by *Eq. 7*. Like the radiation model described in Slatkin (1993), our analytical model represents an instantaneous range expansion in a 2D stepping-stone model. The spatially-structured population is subjected to habitat contraction at time *τ*_1_ in the past (*τ*_1_ < *τ*_2_), during which, as in real study cases, suitable demes are removed from the periphery of the habitat patch. The reduction in *patch-size* has two important consequences expected to influence deme heterozygosity: *i*) decrease in the total amount of habitat, due to loss of suitable habitat at the periphery; and *ii*) increase in edge-influence, caused by the increased proximity of demes to the habitat boundaries. Habitat loss is modelled by considering the total population size before *N*_*τ*_2__*d*_*tot*(*τ*_2_)_ and after *N*_*τ*_1__*d*_*tot*(*τ*_1_)_ contraction. Changes in edge-influence for a given deme *i* are parametrized by two *N_edge_* values, before *N*_*edge*(*i,τ*_2_)_ and after *N*_*edge*(*i,τ*_2_)_ contraction.

Using the cumulative density function (CDF) of probability of identity-by-descent for two alleles sampled from the same deme (Wilkins, 2004), we can then derive the expression for deme heterozygosity at each point in space and time. By rearranging *Eq. 5*, we obtain:

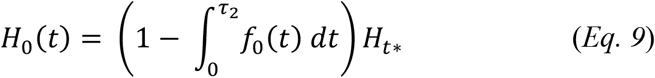

In this expression the ‘separation-of-timescales’ is considered by modelling the ‘*scattering-phase*’ with 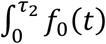 and the ‘*collecting phase*’ with *H_t*_*. The ‘*scattering-phase*’ can be decomposed in two components: the probability of identity-by-descent for two alleles coalescing before *τ*_1_ (time of contraction) and probability of identity-by-descent for two alleles coalescing between *τ*_1_ and *τ*_2_ (time of instantaneous expansion):

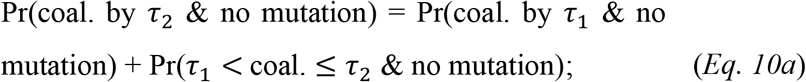

which can be written as:

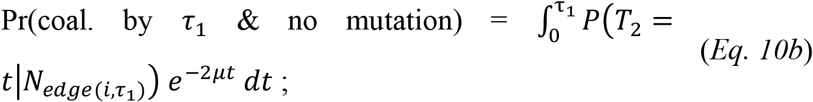

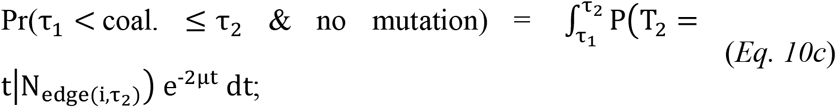

where *P*(*T*_2_ = *t*|*N*_*edge*(*i,τ*_2_)_) or *P*(*T*_2_ = *t*|*N*_*edge*(*i,τ*_1_)_) is the probability of two alleles sampled from the same deme of coalescing at time *t*, given a deme population size of *N*_*edge*(*i,τ*_2_)_ or *N*_*edge*(*i,τ*_1_)_, respectively, before or after habitat contraction; *e*^−2*μt*^ is the probability of no mutation since coalescence time *t*. Then, the CDF of probability of identity-by-descent for two alleles sampled from the same deme at time *t*_0_ is:

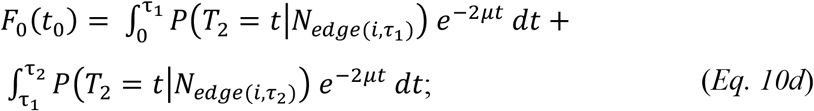

We write the probability that two lineages coalesce at time *t* in the past *P*(*T*_2_ = *t*) using *Equation A7* from Wilkins (2004), which is an approximation of the exact probability distribution in a 2D stepping-stone model (Barton and Wilson 1995, 1996). This approximation allows easier and faster computation while being sufficiently accurate:

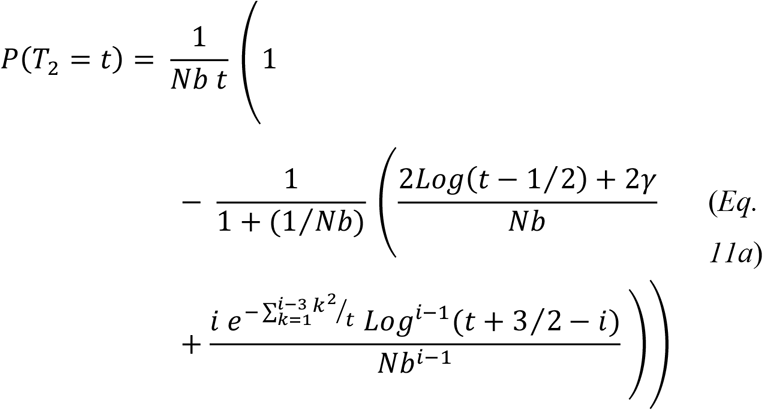

where *γ* is the Euler constant (~ 0.5772), *Nb* is the neighbourhood size (2*N_e_mπ*) and *i* is the number of additional terms for the approximation form, which are added up from three through forty-four with alternate sign. For instance:

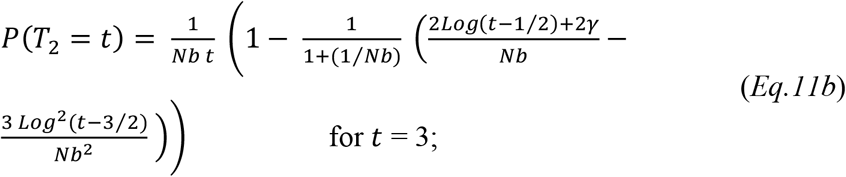

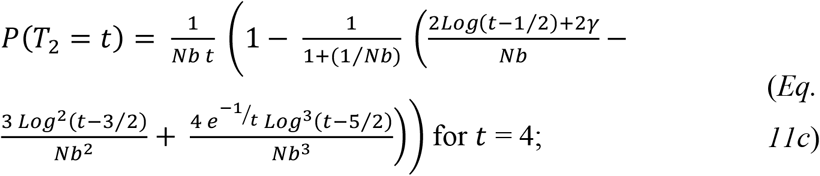

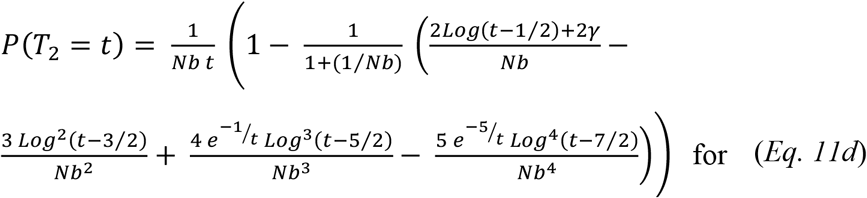

For *t* ≥ 44, all additional terms are included. *Eq. 11* can be substituted in *Eq. 10*, by using either *N*_*edge*(*i,τ*_2_)_) or *N*_*edge*(*i,τ*_2_)_).

Wilkins, (2004) has shown that the ‘*collecting phase*’, under certain conditions, can be approximated by the standard neutral coalescent model (SNCM; Kingman, 1982a, b). As in Barton et al., (2013) and in agreement with the definition of ‘*collecting phase*’ by Wakeley, (1999), *H_t*_* represents the probability that two lineages are heterozygous, independently from geography since these two lineages have most likely crossed the whole habitat patch multiple times. Since we are modelling habitat contraction as a reduction in total patch size and given that the ‘*collecting phase*’ can be approximated by SNCM, *H_t*_* is derived from the expression of the expected heterozygosity for a panmictic population in a varying environment (Griffiths and Tavaré, 1994). In particular, since genetic diversity at time *t* is influenced by past events that decreased significantly the patch population size, we used the harmonic mean of the varying population size over time (Griffiths and Tavaré, 1994) as effective population size from which estimate *H_t*_* by using *Eq. 7*:

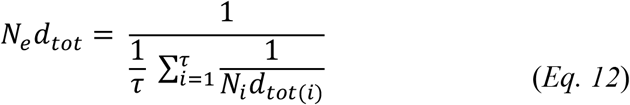

A graphical representation of the non-equilibrium model is presented in Supplementary Appendix S1: Figure S2.

### Population genetics simulations

We used the computer program SINS (Simulating INdividuals In Space; Maié et al., 2021; Rasteiro et al., 2012) to simulate habitat patches undergoing contraction. SINS is an individual-based software that uses a forward-time approach to simulate diploid individuals (males and females) over a 2D grid of demes, similar to a classical 2D stepping-stone model (Kimura and Weiss, 1964) as in SPLATCHE software (Currat et al., 2004; Currat et al., 2019). Deme population size 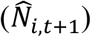 is drawn from a Poisson distribution with a mean value (*N*_*i,t*+1_) given by a corrected form of the Maynard-Smith and Slatkin, (1973) logistic growth (Rasteiro et al., 2012; Maié et al., 2021). The *N*_*i,t*+1_ value in each deme is logistically regulated by deme-specific carrying capacity (*K*), growth rate (*r*) and deme number of reproductive females (*N_f, t_*):

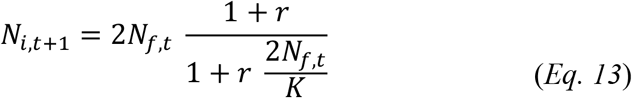

Mating is random within demes, namely one reproductive individual from each sex is randomly chosen and generate a new individual (offspring) which inherit, for each locus, one allele from each parent. At each generation, mating occurs iteratively until 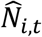 is reached. Each deme exchanges migrants with the four neighbouring demes. The number of individuals that will emigrate from the focal deme at each generation is drawn from a Poisson distribution with mean *M* = *N_i,t_m n_d_*/4, where *m* is the dispersal rate and *n_d_* is the number of available neighbouring demes, which is equal to 2 for a corner deme and 4 for central deme. The number of migrants will then be stochastically distributed among the neighbouring demes using a binomial distribution with probability *P_dir_* = (1 – *F_dir_*)/(*n_d_ – F_t_*), where *dir* is one of the available directions (min=1; max=4), *F_dir_* is a deme-specific friction parameter defining the difficulty to move from the focal deme into that direction and *F_t_* is the sum of the frictions of the *n_d_* receiving demes. SINS is able to simultaneously simulate demographic and genetic data over non-overlapping discrete generations and through space. Several types of molecular markers (sequences, SNPs and microsatellites) and chromosome objects (X and Y chromosomes, autosomes and mitochondrial DNA) can be simulated. In the present study we will exclusively use independent microsatellite loci, which mutations are modelled according to the Stepwise Mutation Model (Ohta and Kimura, 1973). Similar to SPLATCHE (Currat, Ray and Excofier, 2004; Currat et al., 2019), changes from one habitat configuration to another (i.e., habitat contraction) are defined by *K* and *F* maps describing, respectively, the carrying capacity and friction value of each deme in the two configurations. Demes that are unsuitable after an ‘environmental event’ are set with *K* = 0 and *F* = −99, such that no individuals can grow and no migrants can move within these unsuitable demes.

Simulations start with all demes having the same initial genetic diversity, which is defined by a file for each locus specifying the initial number of alleles and frequency of the simulated markers. In this study, we used initial allele frequency for each locus by sampling 100 individuals from a Wright-Fisher (WF) population simulated in *fastsimcoal* (Excoffier and Foll, 2011) using the *fastsimcoal* function in the *strataG* R package (Archer, Adams and Schneiders, 2017). The size of the WF population was set equal to 10,000 and mutation rate according to the value specified in the SINS simulation.

### Simulated scenarios

#### Equilibrium

We simulated mainly two habitat configurations, i) a 9 × 9 squared grid (*d_tot_* = 81 demes) and ii) a 23 × 23 irregular grid (*d_tot_* = 382 demes) (Fig. 1a, b). Three other habitat configurations of 23 × 23 demes were also simulated to show consistency of our results across habitats of different shapes (Supplementary Appendix S1: Figure S6). For each habitat configuration we tested several values of *K* (50, 100, 200, 300) and *m* (0.02, 0.04, 0.06, 0.08, 0.1), which were kept constant across all demes in the habitat. We also tested several values of mutation rate, *μ* (5 × 10^−3^, 5 × 10^−4^, 5 × 10^−5^, 5 × 10^−6^, 5 × 10^−8^) and number of genetic markers (15, 30, 100, 400 microsatellites). The simulations were run for over 10,000 generations, such to ensure that the whole population had reached a ‘quasi-genetic equilibrium’ (i.e., Δ*H_e_/t* ≈ 0). Demographic data (deme population size) were recorded every 50 generations, except when specified otherwise.

**Figure 1.**
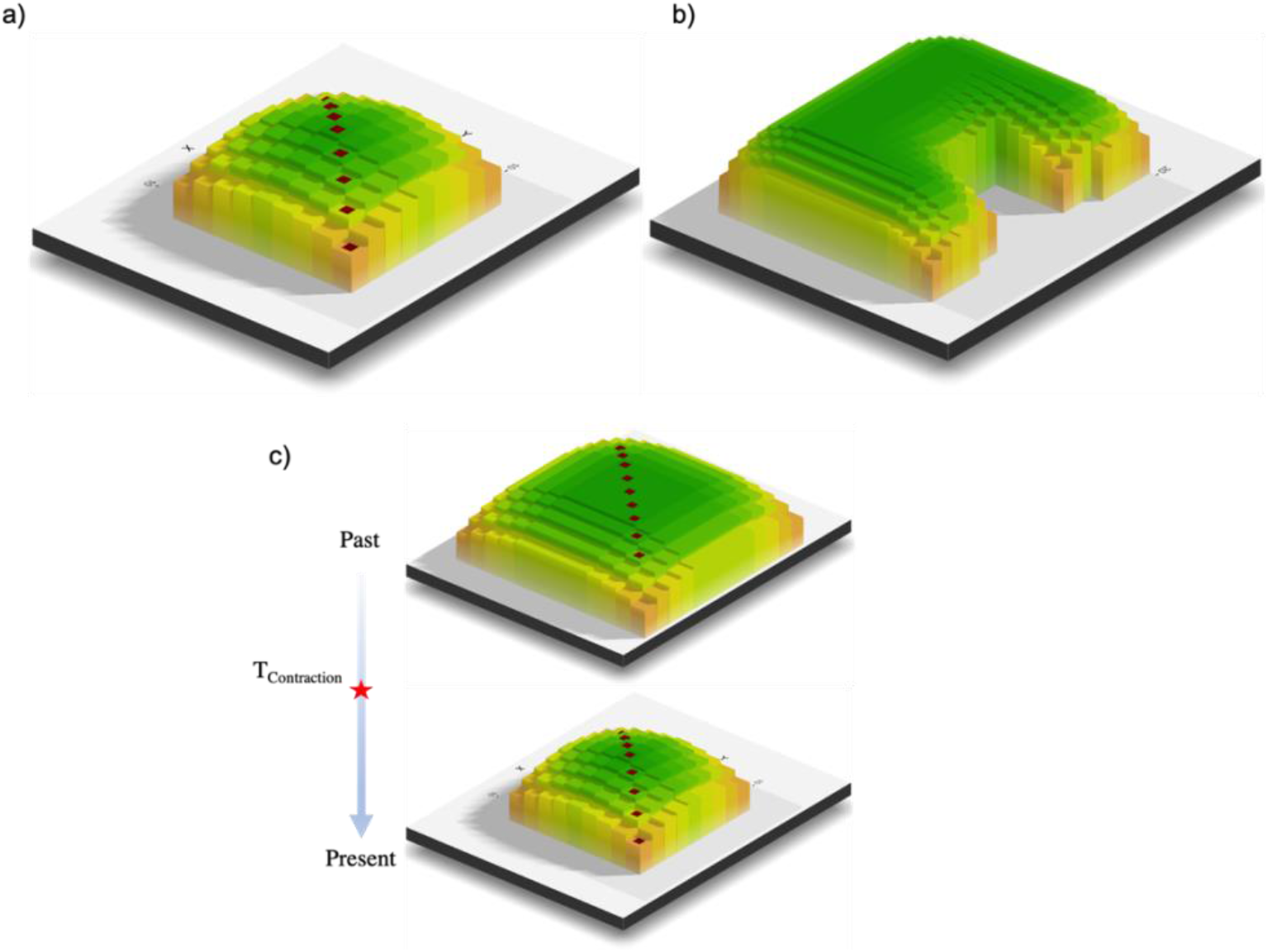
Habitat configurations modelled in this study. **a)** 9 × 9 square grid of demes (d_tot_=81); **b)** 23 × 23 grid of irregular shape (d_tot_=382); **c)** instantaneous habitat contraction from a 13 × 13 (d_tot_=169) to a 9 × 9 grid of demes (d_tot_=81). Within each habitat configuration, demes have homogenous carrying capacity and dispersal rate. Colours and deme bars are proportional to the amount of edge-effect at which each deme is subjected to. Brown squares indicate sampled demes.

#### Non-Equilibrium

We simulated a 13 × 13 squared grid (*d_tot_* = 169 demes) that undergoes habitat contraction after 10,000 generations. Habitat contraction was modelled as an instantaneous removal of the outer demes from the 13 × 13 grid, leading to a 9 × 9 squared grid (Fig. 1c). After habitat contraction, simulations were run for more 4,000 generations. Because of the computational cost of the simulations, most *non-equilibrium* scenarios were simulated only under the 13 × 13 squared grid configuration. However, we also tested the effect of habitat contraction on grid configurations with larger number of demes and more complex shape (Supplementary Appendix S1: Figure S6). As for the *Equilibrium* section, we investigated the impact of *K* (50, 100, 200, 300), *m* (0.02, 0.04, 0.06, 0.08, 0.1) and *μ* (5 × 10^−3^, 5 × 10^−4^, 5 × 10^−5^, 5 × 10^−6^, 5 × 10^−8^) on pattern of genetic diversity, in space and time. Across all *non-equilibrium* simulations, we simulated thirty independent microsatellite loci and sampled fourteen individuals per deme every 50 generations. For both *Equilibrium* and *Non-Equilibrium* settings, we ran 30 independent replicates for each simulated scenario.

### Genetic diversity

Measures of genetic diversity were estimated in R (R Core Team, 2015) using the *summary* function of the *adegenet* R package (Jombart, 2008; Jombart and Ahmed, 2011). We computed three summary statistics typically used in population genetic studies: observed heterozygosity (*H0*), expected heterozygosity (*H_exp_*) and mean number of alleles (*MNA*), which here is equivalent to allelic richness (*Ar*) since sample size is the same across demes (Allendorf and Luikart, 2009). Diversity was estimated for each independent genetic marker and then averaged across loci. Individual-based heterozygosity (*HeL*) was derived by the ‘Homozygosity by loci’ index proposed by Aparicio et al., 2006 and implemented in the *Rhh* R package (Alho et al., 2010). ‘Heterozygosity by loci’ can be computed as *H_e_L* = 1 – *HL* = 1 – ∑ *E_h_*/(∑ *E_h_* + ∑ *E_j_*), where *E_h_* and *E_j_* are the expected heterozygosities of the loci that an individual has in homozygosity (*h*; the two alleles are equal) and in heterozygosity (*j*; the two alleles are different). This index weights locus homozygosity by the level of genetic variability at that locus in the population, such that a locus will contribute more to *HL* if its allele distribution is less skewed (more uniform) and more variable (multiple-alleles). We chose *HL* since it appeared to correlate better with genome-wide homozygosity and inbreeding coefficients in natural populations (Aparicio et al., 2006).

### Demographic analysis

During the spatio-temporal simulation, census population size was recorded for each deme in the habitat. Data were retrieved every 50 generations for the whole simulation, and the harmonic mean of the population size over time was calculated as in *Eq. 12*, but replacing *N_i_d_tot_*(*i*) with *N_i,t_*, the census population size of deme *i* at generation *t*. The harmonic mean of the population size can be used to represent the rate of genetic drift of a given population, when population size varies over time due, for instance, to environmental oscillations (Griffiths and Tavaré, 1994).

## Results

### Equilibrium

We modelled a 2D stepping stone with a total of 81 demes on a 9 × 9 squared grid under homogenous carrying capacity and dispersal rate (Fig. 1a). After 10,000 generations, we sampled 5 individuals from each deme located along the diagonal of the simulated grid. Thirty microsatellites per individuals were ‘genotyped’ and used for estimating deme genetic diversity. In Fig. 2, we show the spatial distribution of deme observed heterozygosity (*H_o_*), assessing the impact of population size, dispersal rate and mutation rate on this measure of genetic diversity. We observed a significant decrease in *H_o_* in demes located on the corner of the habitat (x = 0 and × = 8) compared to central demes (x = 4). Differences in *H_o_* between central and corner demes were stronger in scenarios with low population size (*K* = 50, Δ*H*_0_ = ~0.20; Fig. 2a), low dispersal rate (*m* = 0.02, Δ*H*_0_ = ~0.15; Fig. 2b) and mutation rate (10^−4^ ≤ *μ* ≤ 10^−5^, Δ*H*_0_ = 0.2; Fig. 2c). At these parameter values, we also observed a stronger loss of *H_o_* relative to the maximum level of *H_o_* measured in the habitat (central demes) 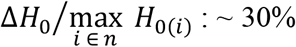 for *K* = 50, ~ 25% for *m* = 0.02 and ~ 30% for *μ* = 10^−4^ (Supplementary Appendix S1: Figure S4). While results on expected heterozygosity are consistent with those on observed heterozygosity, MNA shows rather different patterns (Supplementary Appendix S1: Figure S3, S4). In particular, it is observed that the extent on which central and corner demes differ in diversity is no longer dependent on population size nor on dispersal rate, whereas is still dependent on mutation rate (lower row in Supplementary Appendix S1: Figure S3).

**Figure 2.**
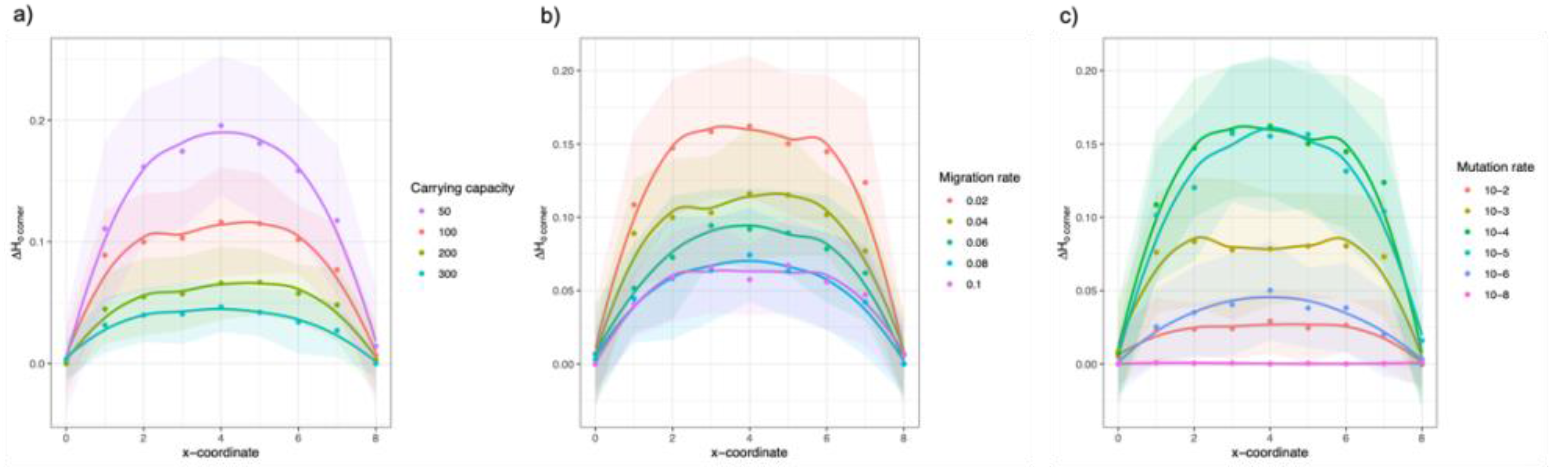
Spatial distribution of differences in observed heterozygosity (*H*_0_) relative to corner demes (Δ*H_o corner_*) along the diagonal of a 9 × 9 stepping-stone model. Impact of **a)** carrying capacity (fixed parameters, *m*: 0.04; *μ*: 5 × 10^−4^); **b)** migration rate (fixed parameters, *K*: 100; *μ*: 5 × 10^−4^); and **c)** mutation rate (fixed parameters, *K*: 100; *m*: 0.02). Central demes (x = 4) show significantly higher *H_o_* with respect to corner demes (x = 0 and × = 8) at lower carrying capacity (*K*: 50; Δ*Ho corner*: ~ 0.20), lower migration rate (*m*: 0.02; Δ*H_o corner_*: ~ 0.15) and intermediate mutation rate (*μ*: 5 × 10^−4^ / 5 × 10^−5^; Δ*H_o corner_*: ~ 0.20).

We tested whether differences in genetic diversity between corner and central demes of the same scenario were attributable to differences in population size (due to demographic stochasticity). Since SINS software simulate population size as a stochastic process, thus allowing some variation in 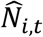 over time, we computed the harmonic mean of the recorded population sizes (HMP) for each deme in the habitat. Supplementary Appendix S1: Figure S5 shows that HMP remain largely unchanged across the whole habitat, an exception being the simulation performed with *K* = 50 and *m* = 0.02, where corner demes experience lower HMP than central demes 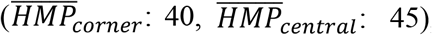, with large fluctuations across simulation replicates (*SD_corner_*: 15, *SD_central_*: 8.5). Note also that HMP was lower than the carrying capacity set at each scenario, approaching to the expected value in simulations with higher carrying capacity.

The results on the 9 × 9 grid provide a concise overview of how edge-effects can influence the spatial distribution of genetic diversity in a finite two-dimensional habitat. Studies that have investigated the influence of proximity to edges on some biological response (ecological or genetic), typically use discrete and simplified definition of edge-effect, such as ‘distance to the nearest-edge’ (see Pfeifer et al., 2017 otherwise). Such approaches do not incorporate the cumulative effect of multiple edges, which is relevant in natural habitat given their common irregular shape. In the present study, we have developed a statistical approach that quantify the additive effect of multiple edges on demes genetic diversity in topologically complex 2D habitat (see *Material and Methods*), similarly to Pfeifer et al., (2017) work, and provided a link with spatial population genetics theory. Using (*Eq. 8*), we assessed the performance of ‘edge-corrected population size’ in predicting the spatial distribution of observed heterozygosity in the habitat of Fig. 1b. In Fig. 3, we compare demes observed heterozygosity with its expected value according to the Wright-Malécot approximation (*Eq. 8*), using as *N_e_* the proposed ‘edge-corrected population size’ (*N_edge_*). As an alternative approach, we perform model-fitting under a GAM model using the ‘edge-corrected population size’ as predictive variable and assess how each independent simulation replicate deviates from the fitted model. Both approaches suggest little deviation between observed and expected values, regardless which population size, dispersal rate or mutation rate is simulated. Results were consistent when simulating other habitat shapes (Supplementary Appendix S1: Figure S6, S7). Note that across all simulated shapes, we observed a strong deviation between theoretical predictions and simulated data when *K*: 50 and *m*: 0.02 (first row, Supplementary Appendix S1: Figure S8). Accordingly, demographic data shows large variability in deme population size at these parameter values, and a lower HMP compared to higher dispersal rates (second row, Supplementary Appendix S1: Figure S8).

**Figure 3.**
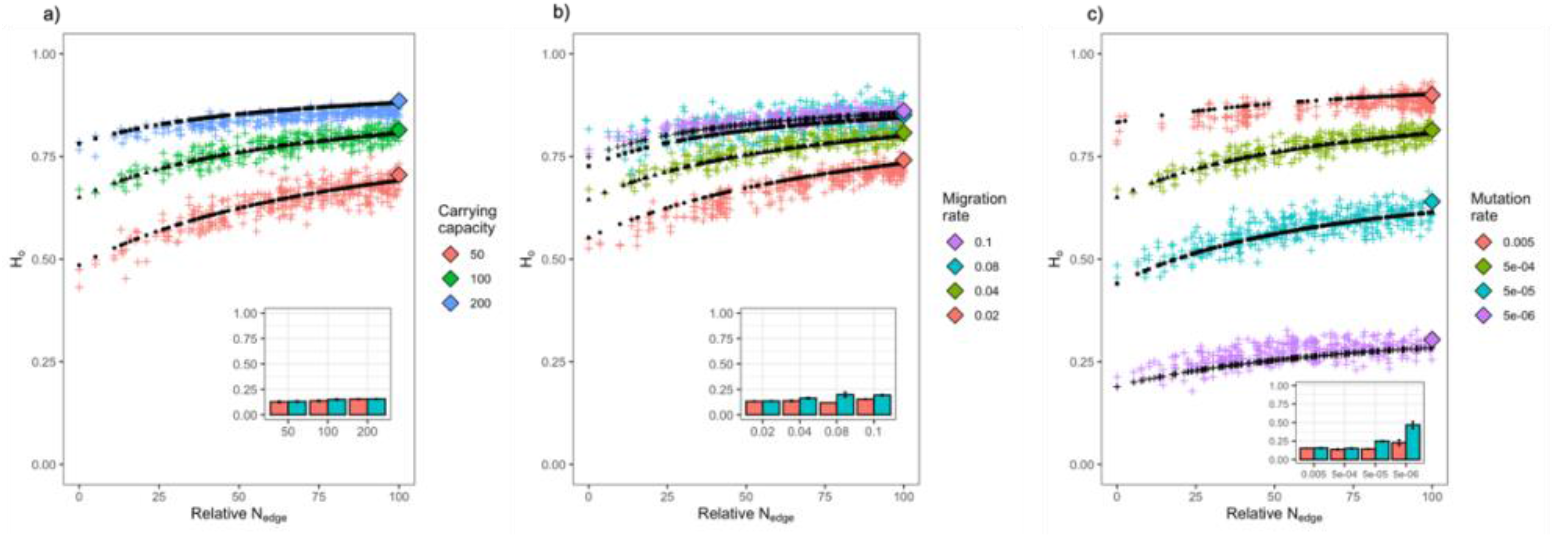
Comparison between edge-corrected theoretical results and simulations on a 23 × 23 stepping-stone model of irregular shape. Impact of **a)** carrying capacity (fixed parameters, *m*: 0.04; *μ*: 5 × 10^−4^); **b)** migration rate (fixed parameters, *K*: 100; *μ*: 5 × 10^−4^); and **c)** mutation rate (fixed parameters, *K*: 100; *m*: 0.04) on the predictability of edge-impacted changes in genetic diversity. Each cross represents the average deme Ho across 30 simulation replicates and 30 microsatellites. Relative N_edge_ is a standardized edge-corrected population size (between 0 and 100) for visualization purposes. Black dots define the theoretical predictions using N_edge_ in the Wright-Malécot approximation (*Eq. 8*). The insert bar plots quantify the NRMSE under the Wright-Malécot approximation (light blue) and GAM model (red). Colored diamonds indicate the expected Ho using the Wright-Malécot approximation and ignoring edge-effect. Under the tested combination of parameters, the results suggest good agreement between the theoretical predictions and simulation results.

One of the parameters that needs to be chosen for quantifying edge-effect is the side length of the area from which habitat information are used to compute the weighted average deme populations size. We show that the side length value can be easily selected and it has little effect on the performance of the statistics (Supplementary Appendix S1: Figure S9). This is because the mean is weighted using a correlation measure (*Eq. 3*) that decays quickly with geographical distance.

We also assessed the effect of the number of genetic markers (15, 30, 100 and 400 loci) on the detectability of edge-impacted changes in population- and individual-based heterozygosity. In Supplementary Appendix S1: Figure S9a-b, we plot the observed heterozygosity of each sampling unit (population or individuals) measured from one simulation replicate with its estimated ‘edge-corrected population size’. We then compare the fitted GAM model from one simulation replicate (coloured line) with i) the fitted GAM model obtained by averaging population or individual-based heterozygosity across simulation replicates (black line); and ii) the theoretical prediction based on the Wright-Malécot approximation (dashed blue line). Overall, the results show a good fit between observed heterozygosity and theoretical prediction (Supplementary Appendix S1: Figure S10c). However, lower number of genetic markers increases the variance around the expected value, and therefore decreases the percentage of genetic variance explained by *N_edge_* (Supplementary Appendix S1: Figure S10d). The effect of the number of genetic markers is stronger with individual-based heterozygosity (9% - 43% variance explained) than with population-based one (27% - 73% variance explained). Since often in real datasets, few individuals are sampled from the same location, we assessed whether binning individual heterozygosity by value of *N_edge_* would improve the percentage of variance explained by the model. The results suggest that individual-binning data analysis improves the explanatory power of *N_edge_* at individual level (from ~9% to ~40% with 15 loci; Supplementary Appendix S1: Figure S11), and therefore could be used when both number of markers and sampled individuals is limited.

### Non-Equilibrium

Since genetic patterns measured from natural populations are unlikely to be at equilibrium, we also investigated the spatio-temporal dynamics of neutral genetic patterns in a habitat undergoing contraction (Fig. 1c). We were particularly interested in exploring the effect of edges on genetic diversity over time, and identify the time required to observe significant differences between central and the edge populations. In Fig. 4 and Supplementary Appendix S1: Figure S12, we show the temporal changes in heterozygosity for central, edge and corner demes, and assess the impact of population size, dispersal rate and mutation rate on these patterns. Overall, we observed that the genetic diversity of corner and edge demes were consistently decreasing over time, with corner demes being more affected than edge demes. Instead, the heterozygosity of central demes did not change much across most parameter combinations of the simulated scenario. However, our simulations show that when dispersal rate is rather high (e.g., *m* > 0.08) or carrying capacity and mutation rate are small (e.g., *K* < 50; *μ* < 10^−4^), central demes are also affected by a decrease in heterozygosity (Fig. 4). The effect of high dispersal rate on central deme genetic diversity is stronger when ‘mean number of alleles’ (MNA) is used as measure of genetic diversity (Supplementary Appendix S1: Figure S13b). Similar to heterozygosity, the MNA of central demes remains unchanged at higher mutation rate (*μ* ≥ 10^−4^), whereas it decreases for lower values of mutation rate (Supplementary Appendix S1: Figure S13c). Unlike dispersal and mutation rate, deme carrying capacity appears to not influence the spatial distribution of MNA and the absolute differences in MNA between central and corner demes (Supplementary Appendix S1: Figure S12a).

**Figure 4.**
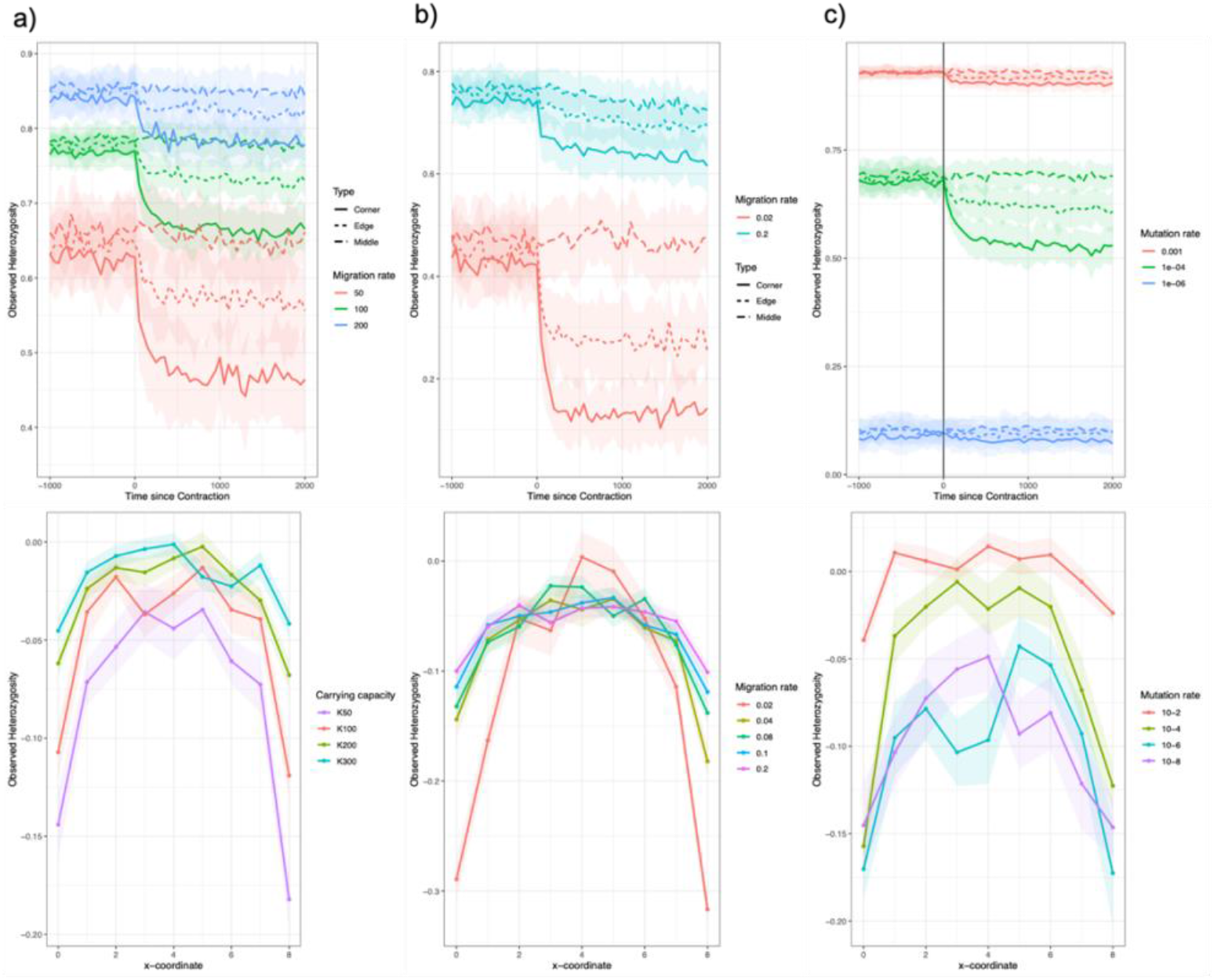
Temporal dynamics of deme *H*_0_ in a 13 × 13 stepping-stone model undergoing habitat contraction. Impact of **a)** carrying capacity (fixed parameters, *m*: 0.04; *μ*: 5 × 10^−4^); **b)** migration rate (fixed parameters, *K*: 100; *μ*: 5 × 10^−4^); and **c)** mutation rate (fixed parameters, *K*: 100; *m*: 0.04) on temporal changes in deme *H*_0_. The first-row shows *H*_0_(*t*) for middle, edge and corner demes. The second row represents the difference in deme *H*_0_ before and after contraction for demes sampled along the diagonal. Across all parameter’s combinations, we sampled 14 individuals per deme, 30 microsatellite per individual. Dots and shade indicate, respectively, the mean and standard deviation of deme *H*_0_ based on 30 simulation replicates.

The simulation results show *i*) how patterns of genetic diversity are affected over time by a decrease in total population size (habitat contraction) and *ii*) by an increase in edge-influence. They provide useful information on the effect of population size, dispersal rate and mutation rate on the temporal dynamic of deme genetic diversity. In addition, we have developed an analytical model that describe how the probability of identity-by-descent of two alleles sampled in the same deme changes over time under habitat contraction (*F*_0_(*t*); see *Material and Methods*). From *F*_0_(*t*), we can derive *H*_0_(*t*), and thus predict how genetic diversity change during habitat contraction. The model considers a 2D stepping-stone model that has been evolving for *τ*_2_ generations, subjected to habitat contraction at *τ*_1_ generations in the past (*τ*_1_ < *τ*_2_). At *τ*_2_ generations in the past, we assume an instantaneous range expansion as in the radiation model described in Slatkin (1993). See Supplementary Appendix S1: Figure S2 for a graphical representation of the analytical model.

Habitat contraction decreases total population size and increases deme edge-influence, affecting the expected pattern of deme *H*_0_(*t*) across the habitat patch. The model implements these processes using four parameters: deme edge-corrected population size before (*N*_*edge*(*i,τ*_2_)_) and after (*N*_*edge*(*i,τ*_1_)_) contraction, and total population size before (*K d*_*tot*(*τ*_2_)_) and after (*K d*_*tot*(*τ*_1_)_) contraction. Assuming that we have prior knowledge on mutation rate, dispersal rate and patch-size, it is only required to estimate *N*_*edge*(*i,τ*_2_)_ and *N*_*edge*(*i,τ*_1_)_ from habitat cover data (e.g., remote-sensing data) using the proposed edge-corrected population size approach (see above, *Material and Methods*’). To describe the dynamic of deme genetic diversity during habitat contraction, we computed numerically *F*_0_(*t*) from *Eq. 10d* and calculated *H*_0_(*t*) from *Eq. 9*. In Fig. 5, we compare simulations and analytical results for three demes with different edge-influence (corner, edge and centre demes) across several values of deme carrying capacity, dispersal and mutation rate. The results refer to the scenario defined in Fig. 1c and suggest good agreement between theoretical *H*_0_(*t*) and simulated deme 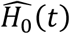. We assessed the theoretical predictions of the analytical model in habitat patches of different shapes and size (Supplementary Appendix S1: Figure S14-S16), randomly sampling fifteen demes across the habitat patches. Except few instances, most demes showed good fit between simulation and analytical results.

**Figure 5.**
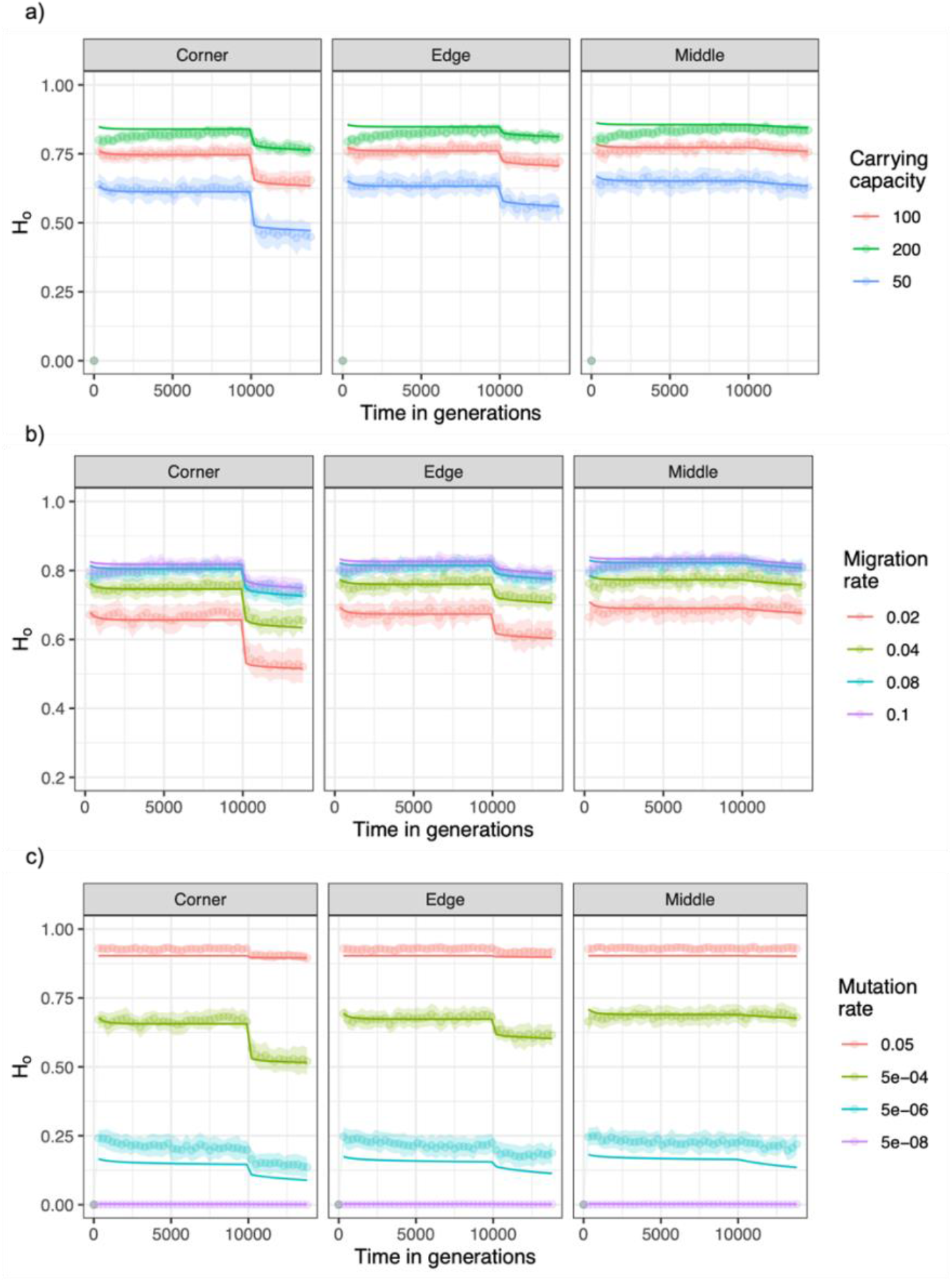
Comparison between analytical and simulation results of a spatially-structured model undergoing habitat contraction. The solid lines represent theoretical predictions based on numerical integration of (*Eq. 10d*). The dots indicate the average deme *H_0_(t*) based on 30 simulation replicates by genotyping 14 individuals per deme at 30 microsatellite loci. We investigated the agreement of analytical and simulation results at various values of **a)** carrying capacity (fixed parameters, *m*: 0.04; *μ*: 5 × 10^−4^), **b)** migration rate (fixed parameters, *K*: 100; *μ*: 5 × 10^−4^) and **c)** mutation rate (fixed parameters, *K*: 50; *m*: 0.04).

## Discussion

In this work we investigated the effect of habitat contraction on the spatial distribution of genetic diversity in species with geographically-limited dispersal. During such process, *habitat-amount* and *edge-effect* play a fundamental role in determining genetic diversity within habitat, and those were the major focus of our work. First, we provided an overview of how edges influence genetic diversity depending on deme population size, migration rate and mutation rate. Second, we proposed a measure called ‘*edge-corrected population size*’ (*N_edge_*) that takes into account population size, migration rate and mutation rate to predict the additive impact of multiple edges on genetic diversity, a pattern expected to be common in natural population. Ultimately, we proposed and explored a theoretical framework describing non-equilibrium deme heterozygosity in populations undergoing habitat contraction, pointing out to the role of *edge-effect* in determining the tempo and mode of changes in genetic diversity. Broadly speaking, the intent of the present study was in part to link some aspects of species ecology with population genetic theory, proposing several approaches that can be used to predict and test how habitat structure affect intra-species genetic diversity.

### Edge-effect in equilibrium populations (habitat patches)

Although often acknowledged, the importance of edge-impacted changes in genetic diversity has been largely overlooked in theoretical and empirical studies. Early works by Maruyama, (1972), Tajima, (1990), have suggested that subpopulations in proximity to habitat edges should hold lower genetic diversity than central subpopulations. However, their conclusions were based on indirect evidence, that is on deductions under the hypothesis that edge-subpopulations would have lower dispersal rate. Later theoretical studies (e.g., Leblois et al., 2006) have shown that observed heterozygosity estimated from samples collected close to habitat boundaries are lower than for samples collected in the centre of the habitat. Similarly, Duforet-Frebourg and Slatkin, (2016) have shown that in a finite 2D stepping-stone model edge-effect increases the correlation of allele frequency between demes compared to an unbounded 2D stepping-stone model. They also noted that edge-effect is less strong at lower migration rate, that is deviations between observed gene correlation in a bounded habitat and expectations under an unbounded model are more limited in space, being significant in close proximity to the habitat boundary and negligible moving towards to centre. Here, we investigated with further details the influence of edge-effect on genetic diversity (deme *H_o_, MNA* and *H_e_*). Our results have shown that deme *H_o_* and *H_e_* are negatively affected by edge-effect, at higher extent in population with low population size or migration rate (*K*: 50; *m*: 0.02;), but also in populations with high values of those demographic parameters (e.g., *K*: 300; *m*: 0.1) (Fig. 2a, b; Supplementary Appendix S1: Figure S3a, b first row). Mutation rate showed a ‘non-linear’ behaviour, since edge-effect was stronger at intermediate values 10^−4^ ≤ *μ* ≤ 10^−5^, but still detectable at higher and lower mutation rate (*μ* < 10^−5^; *μ* > 10^−4^) (Fig. 2c; Supplementary Appendix S1: Figure S3c first row). Mean number of alleles (MNA), showed a rather different behaviour. Edge-effect appeared to impact deme MNA independently from population size and migration rate, with an average diversity loss in corner demes of 1.5 alleles (Supplementary Appendix S1: Figure S3a, b bottom row). Deme MNA showed a linear behaviour in relation to mutation rate, with stronger edge-effect at high mutation rate (Supplementary Appendix S1: Figure S3c bottom row). Overall, our results have shown that corner demes can lose 5% - 33% in heterozygosity (*H_o_* and *H_e_*) and 17% - 33% in allelic diversity compared to central demes (Supplementary Appendix S1: Figure S4), suggesting a strong impact of edges on the number of unique genetic variants (alleles). It is well-established that allelic richness (or MNA) can more easily detect differences in effective population size than heterozygosity (Nei et al., 1975; Allendorf, 1986; Greenbaum et al., 2014; Hoban et al., 2014), since the latter depends on an average of allele frequencies across many alleles (in the case of multi-allelic microsatellites). Corroborating our and previous findings, there is also the recent meta-analysis of Almeida-Rocha et al., (2020), showing that MNA was the statistic detecting the most negative effect of habitat disturbance on genetic response (effect size: −0.65 [−0.80, −0.50]) compared to observed and expected heterozygosity (effect size H_obs_: −0.30 [−0.38, −0.22]; effect size H_exp_: −0.36 [−0.42, −0.30]). Our findings on the effect of edges on allelic diversity are even more relevant if we consider that microsatellite MNA appears to be a better proxy for genome-wide SNP diversity (Ryynänen et al., 2007; Väli et al., 2008; Vilas et al., 2015; Fischer et al., 2017) and a better predictor of long-term adaptation than heterozygosity (Allendorf, 1986; Hughes et al., 2008; Caballero and Garcia-Dorado, 2013). Therefore, when genome-wide molecular markers (genome-wide) are not available for the species of interest, which is often the case for non-model organisms, MNA can be used as a summary statistic to test molecular edge-effects.

### Edge-corrected population size: a proxy of edge-determined changes in genetic diversity

Natural habitat fragments have often irregular shape and heterogenous structure. Simple measures for quantifying edge-effect, such as distance to nearest habitat edge, may not rigorously capture the additive effects of multiple edges, likely occurring in natural habitats. This was the central idea behind the ‘edge-influence’ statistics developed by Pfeifer et al., (2017), which they used to assess the impact of multiple additive habitat edges on species abundance across 1,673 vertebrate taxa. Their work was preceded by a global analysis of habitat fragmentation across the world, showing that nearly 70% of remaining forests today are at 1 km of an edge (Haddad et al., 2015), and therefore pointing out the likely pervasive impact of habitat edges on biodiversity. Pfeifer et al., (2017) and other previous studies have investigated the ecological response of species to habitat edges, and currently little is known on edge-impacted changes in genetic diversity on natural populations. Reasons for this knowledge gap are probably related to the necessity of a spatially-restricted sampling design, which may not correspond with the primary objectives of most conservation genetic studies, often looking at *landscape*-scale genetic pattern (DiLeo and Wagner, 2016), but also to the need of a large number of genetic markers for reliably capture fine-scale genetic variation. Few studies, to our knowledge, have explicitly investigated molecular edge-effect in wild populations, namely Radespiel et al., (2019) and Quevedo et al., (2013), although suitable datasets to address this question are likely available in literature.

In this study, we have proposed a framework that uses information on local carrying capacity, species dispersal rate and mutation rate to estimate the additive effect of multiple edges on genetic diversity 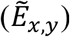. Such measure is then used to correct local carrying capacity (*N_edge_*), and predict edge-impacted changes in heterozygosity by implementing *N_edge_* in the Malécot-Wright approximation for the probability of identity-by-descent of two alleles sampled from the same location (*F*_0(*t*=∞)_ in *Eq. 6*). Our results have shown that *N_edge_* can be a good predictor of spatial genetic variation in finite habitats, under various parameters’ combinations (population size, dispersal and mutation rate) (Fig. 3; Supplementary Appendix S1: Figure S7). The accuracy of the theoretical predictions decreased significantly at lower migration rate (*K*: 50; *m*: 0.02; Supplementary Appendix S1: Figure S8), probably due to the large fluctuations in population size over time at this parameters’ combination (Supplementary Appendix S1: Figure S5a or S7). Another explanation could be given by the behaviour of the Malécot-Wright approximation for two alleles sampled from the same location. In fact, it has been shown that Malécot-Wright approximation deviates significantly from exact results when neighbourhood size (2*Nmπ*) is smaller than 10 (Wilkins, 2004). An alternative explanation refers to the observation of Wilkins, (2004) showing that, when 2*Nmπ* < ~12.5, the genealogical process at the collecting phase can significantly deviate from the standard coalescent. Nevertheless, this parameters’ combination did not have an effect on the fitted GAM model, suggesting that when observed data deviates significantly from theoretical predictions, the *N_edge_* can still be used as explanatory variable. The analytical model performed well also when observed heterozygosity was measured at the individual level (‘Heterozygosity-by-loci’; Aparicio et al., 2006), although, with a lower explanatory power than deme observed heterozygosity (Supplementary Appendix S1: Figure S10c, d). In both types of sampling unit, the number of genotyped markers had a strong impact on the percentage of variance explained by the model, with values of 9% and 27% with 15 loci, and 43% and 73% with 400 loci, respectively at the individual and deme-level (Supplementary Appendix S1: Figure S10d). Binning individual heterozygosity by value of *N_edge_* increases the explanatory power of *N_edge_* on individual based heterozygosity (Supplementary Appendix S1: Figure S11), and thus could be used when sample size is limited in real study cases.

The ‘edge-corrected population size’ proposed in the present study could also be interpreted as a measure of connectivity. As detailed in *Material and Methods*, *N_edge_* is estimated by using the weighted mean of deme carrying capacity within an area (*A*) centred at the focal point. The weight for a deme *i* included in *A* is proportional to the expected genetic correlation between the focal point and deme *i*. The underlying assumption is that deme effective population size at each location in space depends on the census deme-population size, but also on the surrounding amount of habitat from which a given deme can receive migrants at each generation (connectivity). Connectivity indices have a long history in spatial ecology (Moilanen and Nieminen, 2002; Balkenhol et al., 2009; Pfluger and Balkenhol, 2014). In landscape genetics, connectivity measures from spatial ecology have been used to link environmental variables with neutral and adaptive genetic variation (e.g., Keyghobadi et al., 2005; Taylor and Hoffman, 2014). A class of those measures are called ‘neighborhood-based’ approaches which, similarly to our framework, weight the influence of habitat variables by geographical and dispersal distance. However, although spatial ecology and genetics share fundamental mechanisms, they also differ in some processes. In particular, population genetic theory predicts that neutral genetic variation fluctuates over spatial scales of 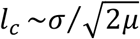 (or 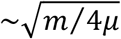 for a 2D stepping-stone model) (Malécot, 1948; Barton et al., 2002), which depends on the balance between dispersal and mutation rate. Since *N_edge_* is measured by explicitly taking into account the ‘spatial scale of variation’ (*l_c_*) of the focal species (*Eq. 3*), it is expected to be a better predictor of genetic diversity than measures of ‘ecological’ connectivity.

In literature, the necessity of a landscape genetic theory that unifies *“ecology and evolution in a truly interdisciplinary way”* has been stated in several occasions (Dyer, 2015; DiLeo and Wagner, 2016; Balkenhol et al., 2015, 2017). With the present work, we aimed to partially contribute to this knowledge gap and, in addition, provide a null-model at equilibrium for molecular edge effect, which to our knowledge has never been proposed.

### Non-Equilibrium pattern of genetic diversity during habitat contraction

The results discussed above make the important assumption that populations are at ‘quasi-equilibrium’, namely that genetic variation remain nearly unchanged over time, representing the local environmental conditions of the population as we see it today. This might be a robust assumption if, as pointed out by Slatkin and Barton, (1989) and Barton et al. (2013), samples are collected over intermediate spatial scale (area A), such that Diameter(A)^2^/*σ*^2^ ≪ *t^*^* ≪ *μ*^−1^, where t*can be interpreted as a time window over which we are sufficiently confident that habitat structure has remained unchanged. However, predictions under the ‘quasi-equilibrium’ assumption do not provide, by definition, an understanding of the temporal dynamics of genetic variation.

Here, we have considered a non-equilibrium spatial model where a geographically-structured population undergoes habitat contraction at a given point in time. First, we have explored through spatial simulations the impact of carrying capacity, dispersal rate and mutation rate on the dynamic of genetic diversity. Our results have shown that genetic diversity is influenced by changes in *patch-size* and *edge-effect* relatively quickly, within few hundreds of generations since contraction. The joint effect of habitat reduction and *edge-effect* on genetic diversity were observed across all parameters’ combinations, including large carrying capacity, high migration rate and high mutation rate. Subpopulations in proximity to habitat edges were the most affected by the contraction event, whereas central subpopulations remained nearly unaffected. This is true for most parameters’ combination except when dispersal rate is high (*m* > 0.02) and mutation rate is low (*μ* ≤ 10^−6^). In fact, under these ranges, central demes showed a decrease in genetic diversity. As observed at *equilibrium* condition, MNA is the measure showing the largest changes in genetic diversity, even for central subpopulations (Supplementary Appendix S1: Figure S13b). Although initially surprising, the larger impact of edges on central demes for highly dispersing populations is deducible from the theory. In fact, the Malécot-Wright approximation includes the quantity *l_c_*, the ‘ spatial scale of genetic variation’, which is proportional to the dispersal rate of the studied species. This implies that changes in neutral allele frequency at a given point in space can fluctuate over relatively large distances in high dispersing populations. These results suggest also that species with high dispersal may in principle be more affected by edge-effect than geographically-restricted populations. Similar observations were reported in Duforet-Frebourg and Slatkin, (2016), showing that edge-effect increases the correlation of allele frequency between demes, at larger extent with high than low migration rate.

In our work, we have in addition developed a theoretical framework describing and predicting the non-equilibrium dynamic of deme genetic diversity following changes in *patch-size* and *edge-effect*. We have shown good agreement between simulation and analytical results, suggesting that such framework could be used for predicting the effect of habitat disturbance on genetic diversity. One of the aspects to stress about our spatial model is that both *patch*-level processes (*edge-effect* and *patch-size*) plays a fundamental role in determining spatial genetic diversity. *Edge-effect* influences deme heterozygosity relatively fast after contraction, depending on deme position relative to edges (Fig. 5; Supplementary Appendix S1: Figure S14-S16). Instead, reduction in *patch-size* impacts deme heterozygosity indistinctly across demes and at a slower rate than *edge-effect*. This reflects the separation-of-timescales in the genealogy of structured models, characterized by a short-term (*scattering-phase*) and a long-term (*collecting phase*) coalescent process (Wakeley, 1999; Wilkins, 2004). We have shown that the short-term process can be modelled using the estimated edge-corrected population size (*N_edge_*), whereas the long-term process can be modelled as the harmonic mean of the total population size (*patch-size*) over time. A similar theoretical problem has been addressed by Alcala et al., (2013), where they analytically characterized the dynamics of genetic diversity following a change in migration rate between populations in a finite island-model. Ours and Alcala et al., (2013) results on within-population genetic diversity are qualitatively concordant. That is, in both structured models, within-population genetic diversity quickly decays over time following an exponential distribution. The rate of such decay is decomposed in two temporal dynamics, a short-term dynamic dependent on *μ, m*, local *N* and total population size; and a long-term dynamic dependent only on *μ* and total population size. However, here we take into account an explicit spatial model (2D stepping stone), since many species are indeed spatially structured (Sexton et al., 2014).

The non-equilibrium framework here proposed has been developed to investigate and predict the effect of habitat contraction (*edge-effect* and *patch-size*) on genetic diversity. In this framework, changes in genetic diversity have been modelled as the results of changes in local and total effective population size. Therefore, we speculate that our non-equilibrium model could be adapted to investigate recent changes in population size in geographically-structured populations. Several approaches have been developed to model changes in population size or/and connectivity in structured models, although often ignoring the effect of space (e.g., Nielsen and Wakeley 2001; Hey and Nielsen 2004; Gutenkunst et al. 2009; Costa and Wilkinson-Herbots 2016; Arredondo et al., 2021). On the other hand, multiple studies have investigated non-equilibrium genetic patterns in homogenous geographically-structured populations (Nagylaki 1974, 1977; Slatkin, 1991; Slatkin, 1993; Duforet-Frebourg and Slatkin, 2016; Ortega-Del Vecchyo and Slatkin, 2019), but to our knowledge not much has been done on non-equilibrium spatial models in varying environments (e.g., habitat contraction).

### Possible applications and future implementations

The ‘edge-corrected population size’ statistic described in the present study has been thought as an approach to test edge-impacted changes in genetic diversity. The equilibrium and non-equilibrium frameworks rely on information that are not necessarily available, but for which sufficiently good proxy can be found. For instance, information of patch habitat quality can be retrieved from tree cover datasets (e.g., Hansen et al., 2013) or from Normalized Difference Vegetation Index (NDVI) (e.g., https://earthdata.nasa.gov/lance). Further extension of our work could include the exploration and validation of an additional statistic quantifying the ‘genetic sensitivity’ of species to edge-effects (see Pfeifer et al., 2017 for details). This would require genetic data across several patches, with sufficient variability in carrying capacity (habitat amount) and ‘genetic edge-influence’. Moreover, estimates of ‘edge-corrected population size’ could provide some insights on how (and if) edge-impacted genetic diversity influences the success of local adaptation in perturbed environments that may arise from the creation of habitat edges. Accordingly, Polechova, (2018), has shown that the success for range expansion or colonization of new environments in a spatially-restricted species is strongly dependent on genetic drift. In particular, Polechova, (2018) define an ‘expansion threshold’ below which adaptation cannot occur due to the strength of genetic drift, given that it reduces genetic diversity below that required for adaptation to a heterogeneous environment.

Other applications of the ‘edge-corrected population size’ can be imagined. For instance, our approach could be used to correct measure of pairwise genetic differentiation by controlling for the increased genetic drift due to edge-effect. In fact, pairwise *F_st_* computed including edge-populations are significantly higher than pairwise comparisons with no edge populations (Supplementary Appendix S1: Figure S17, S18). Lastly, further exploration of the non-equilibrium model could reveal whether this could be used in an inferential framework to estimate past changes in population size in geographically-structured populations. At this stage, this is largely speculative and would require further investigation.

## Author Contributions

GMS, LC and TM designed the project. GMS performed spatial genetic simulations and data analysis, and developed the numerical framework. TZ contributed to data analysis. TM and RR developed the genetic simulator. GMS drafted the manuscript and LC revised it.

## Acknowledgments

This work was supported by the 2015–2016 BiodivERsA COFUND call for research proposals, with the national funders Agence Nationale de la Recherche (grant number ANR-16-EBI3–0014), Fundação para a Ciência e Tecnologia (grant number Biodiversa/0003/2015) and German Bundesministerium für Bildung und Forschung (grant number 01LC1617A). This work was also supported by the Fundação para a Ciência e Tecnologia (grant numbers PTDC/BIA-BEC/100176/2008, PTDC/BIA-BIC/4476/2012, PTDC-BIA-EVL/30815/2017 to L.C., PD/BD/114343/2016 to G.M.S), by the Laboratoire d’Excellence (LABEX) entitled TULIP (grant numbers ANR-10-LABX-41 and ANR-11-IDEX-0002-02) as well as the the LIA BEEG-B (Laboratoire International Associé–Bioinformatics, Ecology, Evolution, Genomics and Behaviour) and the Investissement d’Avenir grant of the Agence Nationale de la Recherche (grant number CEBA: ANR-10-LABX-25-01).

