## Supplementary Appendix S1 for "On the genetic consequences of habitat contraction: edge effects and habitat loss"

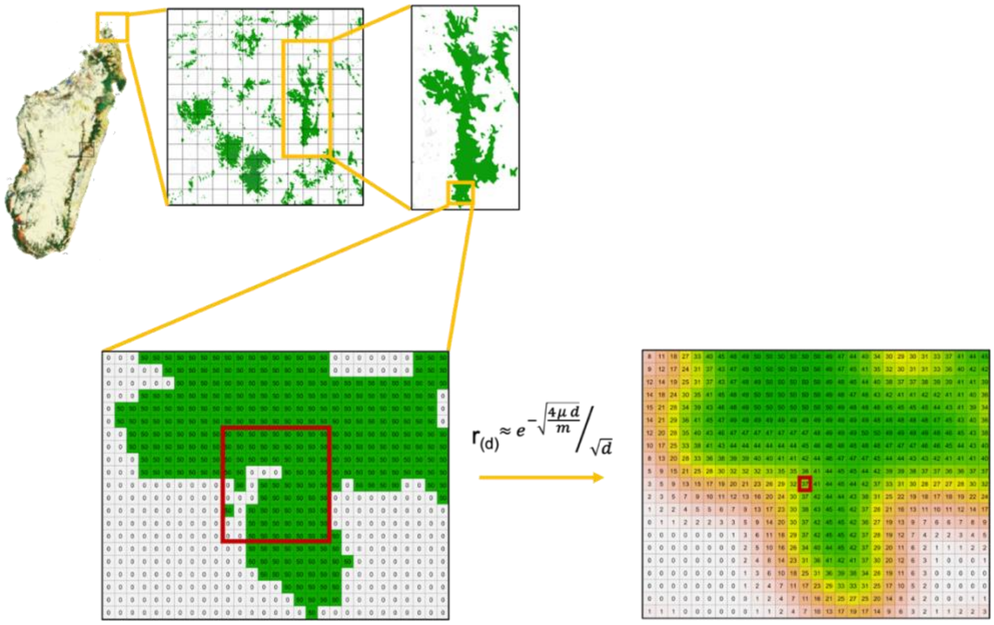


**Figure S1. Illustration of the spatial filtering for computing 'edge-influence index'.** As an example, it is shown the percentage of tree cover for several forest fragments in the Loky-Manambato region, northern Madagascar. We removed detailed information on tree cover by setting the cells with tree cover < 60 to 0, and to 50 otherwise. The solid red line shows a square matrix with area $A$ and side length $R$ = 9. The focal point (in the centre of the square matrix) will get the weighted average of the cells within $A$. The weights are calculated using the genetic correlation equation from Kimura and Weiss, 1964. The genetic correlation equation show that the focal point has the heaviest weight and other cells within A have smaller weights as their distance from the focal point increases.

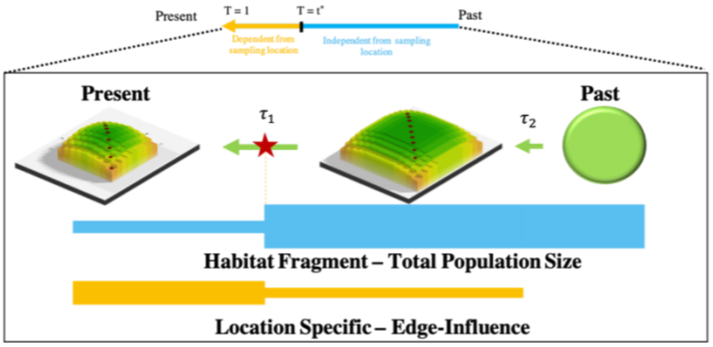


$$H_0(t) = \left(1 - \int_0^{\tau_2} f_0(t)\,dt\right) H_{t*}$$

$$\int_0^{\tau_2} f_0(t)\,dt = \Pr(\text{coal. by } \tau_2 \text{ \& no mutation}) = \Pr(\text{coal. by } \tau_1 \text{ \& no mutation}) + \Pr(\tau_1 < \text{coal.} \leq \tau_2 \text{ \& no mutation})$$

**Figure S2. Illustration of the non-equilibrium spatial analytical model of habitat contraction.** The genealogical process of a spatially structured model can be divided in two phases: *scattering phase* (dependent on alleles sampling location) and *collecting phase* (mostly independent on sampling location). The transition time between these two phases is indicated by $t^*$. Thinking backward in time, two alleles are sampled within subpopulation in the present ($t = 0$). The habitat fragment has undergone contraction at $\tau_1$ and an instantaneous range expansion at $\tau_2$. The analytical model assumes that edge effect influences the *scattering phase*, whereas habitat loss influences the *collecting phase* of the genealogical process.

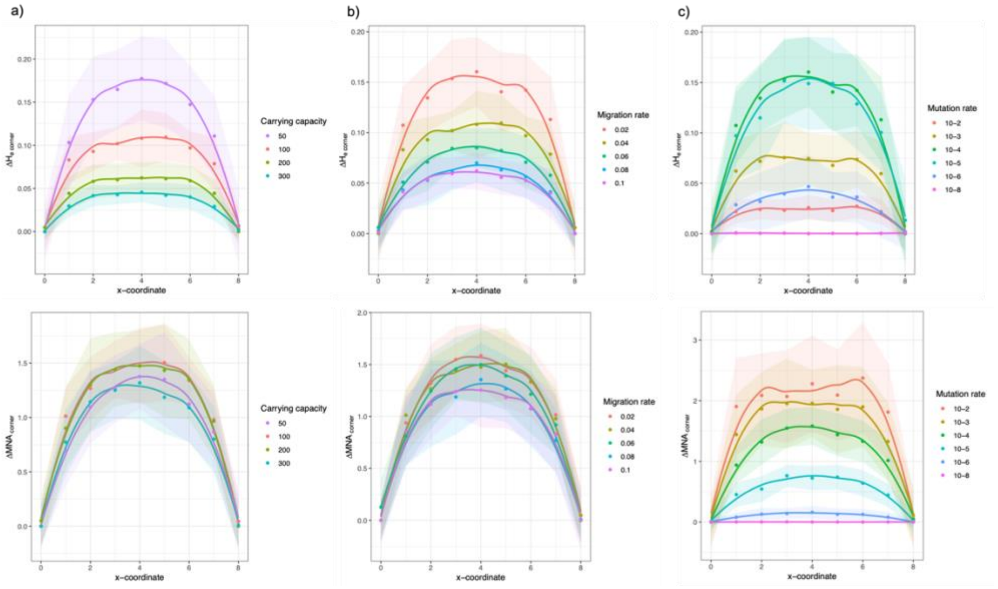


**Figure S3. Spatial distribution of differences in expected heterozygosity ($\Delta H_{e\ corner}$) and mean number of alleles ($\Delta MNA_{corner}$) relative to corner demes along the diagonal of a 9 x 9 stepping-stone model.** Impact of **a)** carrying capacity (fixed parameters, $m$: 0.04; $\mu$: 5 x $10^{-4}$); **b)** migration rate (fixed parameters, $K$: 100; $\mu$: 5 x $10^{-4}$); and **c)** mutation rate (fixed parameters, $K$: 100; $m$: 0.02). Central demes (x = 4) show significantly higher $H_e$ with respect to corner demes (x = 0 and x = 8) at lower carrying capacity ($K$: 50; $\Delta H_{o\ corner}$: ~ 0.17), lower migration rate ($m$: 0.02; $\Delta H_{o\ corner}$: ~ 0.15) and intermediate mutation rate ($\mu$: 5 x $10^{-4}$ / 5 x $10^{-5}$; $\Delta H_{o\ corner}$: ~ 0.15). $\Delta MNA_{corner}$ does not change significantly with carrying capacity and migration rate, whereas it is more pronounced at higher mutation rate (e.g., $\mu$: 5 x $10^{-2}$; $\Delta MNA_{corner}$: ~ 2.2).

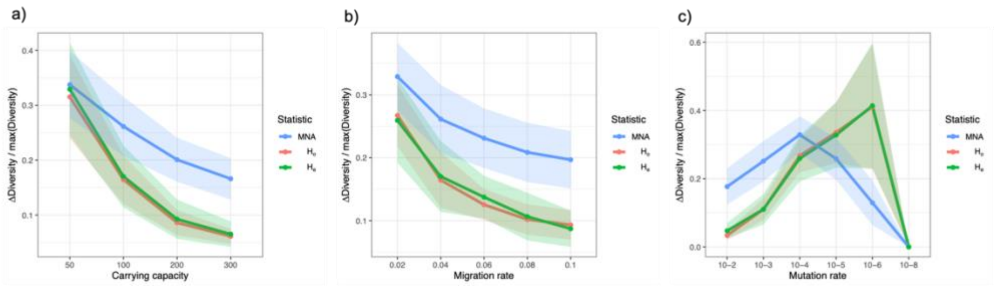


**Figure S4. Relative difference in genetic diversity ($\Delta MNA_{corner}$, $\Delta H_{o\ corner}$, $\Delta H_{e\ corner}$) between central and corner demes sampled in a 9 x 9 stepping-stone model.** Corner demes shows a decrease in genetic diversity of **a)** ~33% or ~5-17% at, respectively, $K$=50 and $K$=300 (fixed parameters, $m$: 0.04; $\mu$: 5 x $10^{-4}$); **b)** ~25-33% or ~10-20% at, respectively, $m$: 0.02 and $m$: 0.1 (fixed parameters, $K$: 100; $\mu$: 5 x $10^{-4}$); and **c)** ~5-20% or ~17-40% at, respectively, $\mu$: 5 x $10^{-2}$ and $\mu$: 5 x $10^{-6}$ (fixed parameters, $K$: 100; $m$: 0.02). $\Delta H_{o\ corner}$ and $\Delta H_{e\ corner}$ increases linearly with decreasing mutation rate up to $\mu$: 5 x $10^{-8}$, since no genetic diversity was present at this value of mutation rate. $\Delta MNA_{corner}$ shows a non-linear effect with respect to mutation rate, being more affected by edge-effect at intermediate mutation rate ($10^{-3}$-$10^{-5}$).

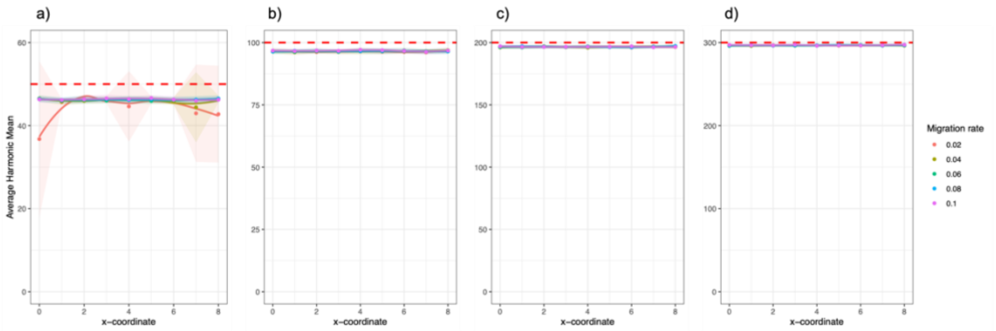


**Figure S5. Harmonic mean of deme population size (HMP) across 10,000 generations in a 9 x 9 stepping-stone model for several values of migration rate. a)** $K$: 50; **b)** $K$: 100; **c)** $K$: 200; **d)** $K$: 300. Dashed red line indicates the simulated carrying capacity for each scenario. Thick lines represent the average HMP across 30 demography simulations, with standard deviation given by the shade around the mean. Note that the harmonic mean was always lower than the value of carrying capacity set at each scenario, with larger deviation for smaller values of carrying capacity. The results do not show differences in HMP along diagonal demes, with the exception of $K$: 50, $m$: 0.02 scenario where corner demes show lower HMP than central demes.

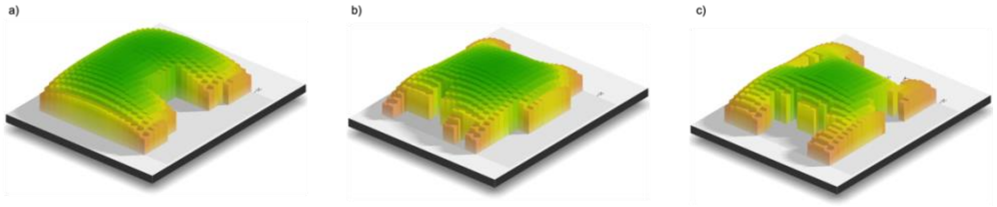


**Figure S6. Simulated scenario for the edge-corrected population size. a)** ‘scenario 1’ including 382 demes, also present in main text; **b)** ‘scenario 2’ including 318 demes; **c)** ‘scenario 3’ including 305 demes. Colors and bars refer to deme-specific edge influence, that for visualization purposes was estimated assuming $K$: 50, $m$: 0.04 and $\mu$: 5 x $10^{-4}$.

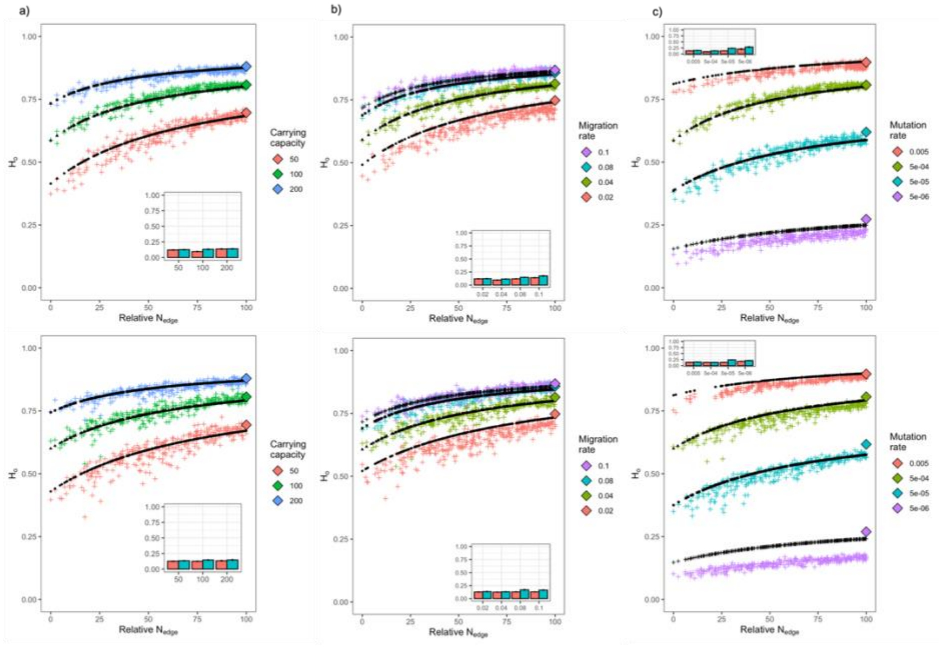


**Figure S7. Comparison between edge-corrected theoretical results and simulations on two 23 x 23 stepping-stone model of irregular shape.** Impact of **a)** carrying capacity (fixed parameters, $m$: 0.04; $\mu$: 5 x $10^{-4}$); **b)** migration rate (fixed parameters, $K$: 100; $\mu$: 5 x $10^{-4}$); and **c)** mutation rate (fixed parameters, $K$: 100; $m$: 0.04) on the predictability of edge-impacted changes in genetic diversity. Each cross represents the average deme $H_o$ across 30 simulation replicates. Relative $N_{edge}$ is a standardized edge-corrected population size (between 0 and 100) for visualization purposes. Black symbols define the theoretical predictions using $N_{edge}$ in the Wright-Malécot approximation (Eq. 6). The insert bar plots quantify the NRMSE under the Wright-Malécot approximation (light blue) and GAM model (red). Colored diamonds indicate the expected $H_o$ using the Wright-Malécot approximation and ignoring edge-effect.

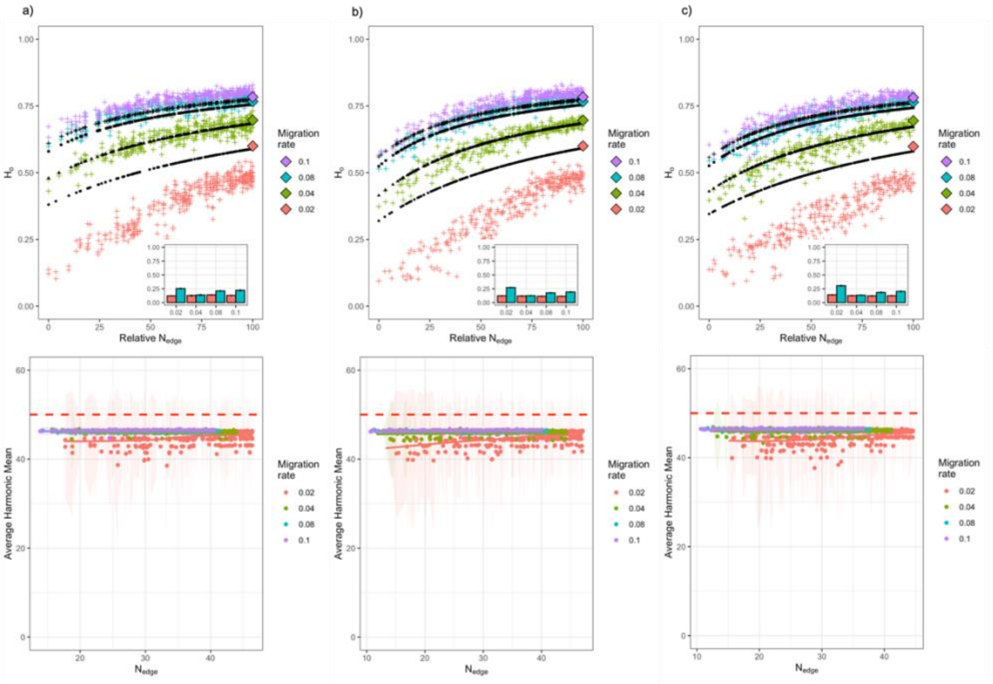


**Figure S8. Comparison between edge-corrected theoretical results and simulations on three 23 x 23 stepping-stone model of irregular shape with $K$: 50.** First row: the effect of varying migration rate on the observed heterozygosity predicted for **a)** scenario 1; **b)** scenario 2; and **c)** scenario 3. See Fig. S5 for a visual representation of the simulated habitat shapes. The insert bar plots quantify the NRMSE under the Wright-Malécot approximation (light blue) and GAM model (red). Second row: harmonic mean of deme population size (HMP) across 10,000 generations with respect to $N_{edge}$. Dashed red line indicates $K$: 50. Standard deviation is given by the shade around the mean. $K$: 50 and $m$: 0.02 shows large variability and lower HMP than higher migration rate, which can explain the poor predictability of $N_{edge}$ at these parameter values.

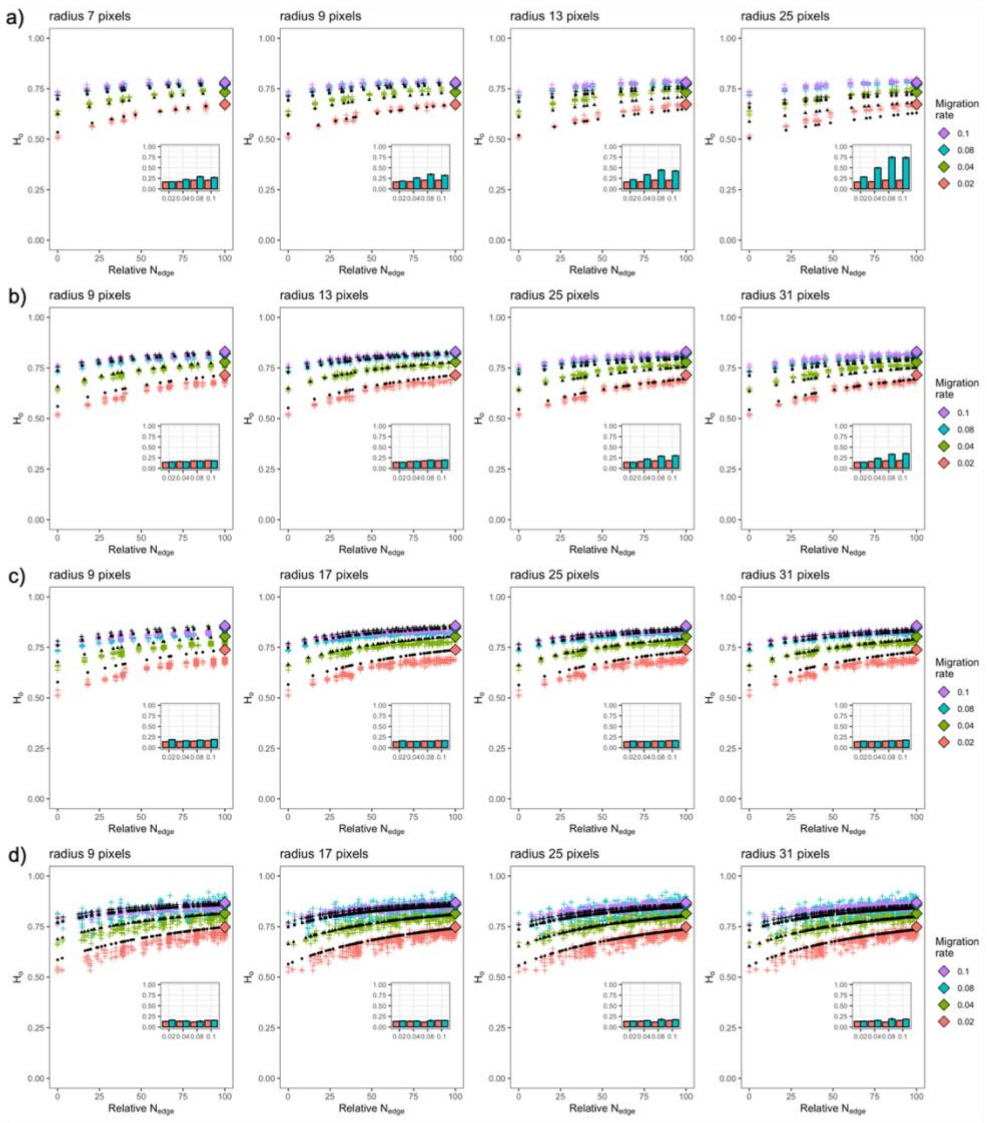


**Figure S9. Effect of radius selection on the edge-corrected population size ($N_{edge}$).** These results compare theoretical expectations and simulations for **a)** 9 x 9; **b)** 13 x 13; **c)** 17 x 17 square grids; and **d)** 'scenario 1' from Fig. S1 under different migration rate values and pixel size for the radius selection (fixed parameters, $K$: 100 and $\mu$: 5 x $10^{-4}$). Each cross represents the average deme $H_o$ across 30 simulation replicates. Colored diamonds indicate the expected $H_o$ using the Wright-Malécot approximation and ignoring edge-effect. The insert bar plots quantify the NRMSE under the Wright-Malécot approximation (light blue) and GAM model (red). We conclude that the radius size has little effect on the performance of $N_{edge}$.

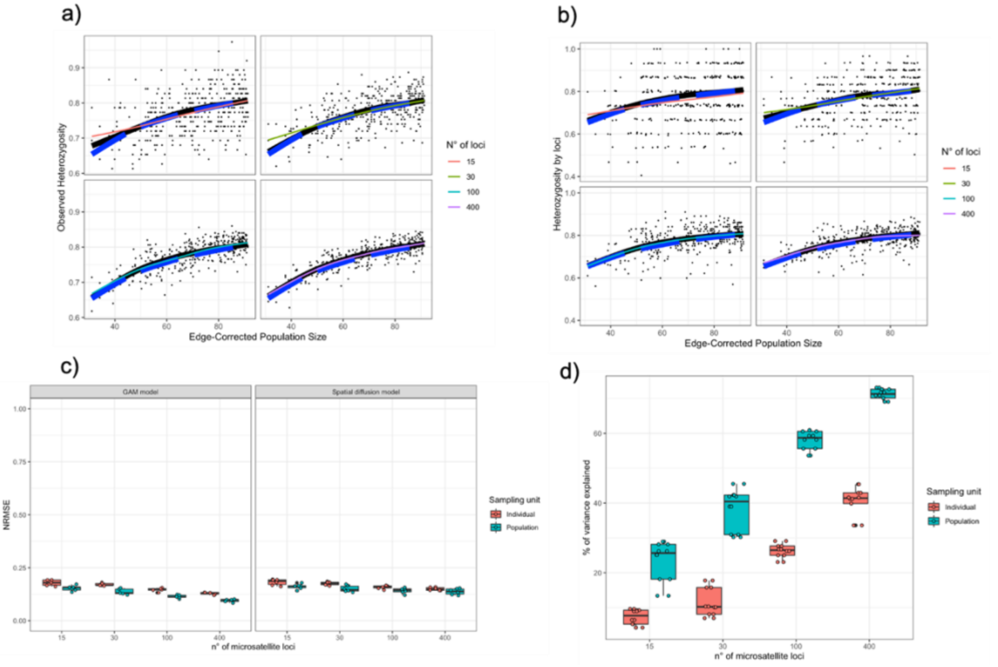


**Figure S10. Effect of the number of genetic markers and sampling unit (deme or individual) on the detection of edge-determined changes in genetic diversity.** The impact of 15, 30, 100 or 400 microsatellite markers on the agreement between theoretical and simulation results for **a)** deme $H_o$ and **b)** individual 'Heterozygosity by loci'. Each point represents genetic diversity estimated from samples collected from one deme in one simulation replicate. Dashed blue and solid black line indicate, respectively, theoretical results and average value across 30 simulations. Colored line is a GAM model obtained from one simulation replicate. **c)** NRMSE and **d)** Percentage of variance explained by the 'edge-corrected population size' for deme $H_o$ (light blue) and individual 'Heterozygosity by loci' (red) when varying number of genetic markers. The results suggest good agreement between theoretical and estimates of genetic diversity from one simulation replicate (as proxy of what can be obtained from field samples). The power of detection is greatly affected by the number of microsatellite markers, if below 100.

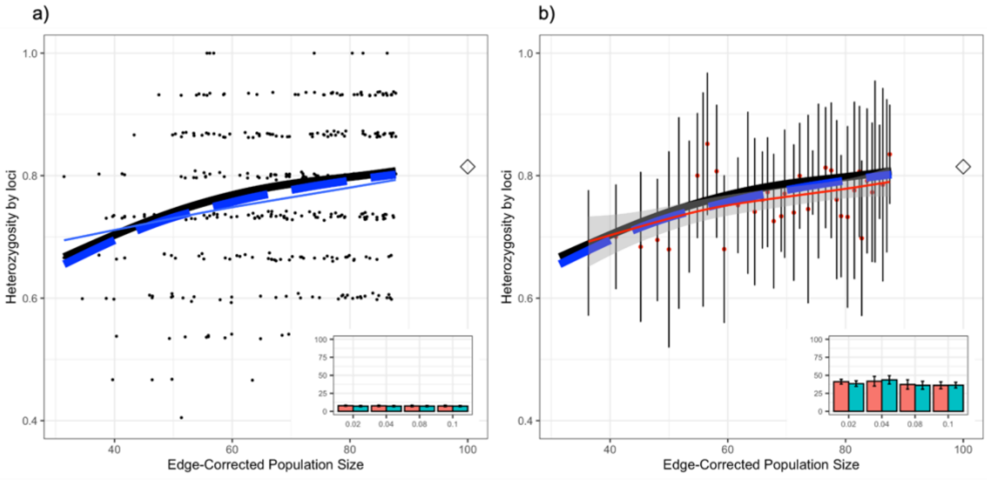


**Figure S11. Effect of edge-corrected population size data binning on individual-based heterozygosity. a)** raw individual-based heterozygosity; **b)** individual heterozygosity grouped in 40 bins of equal sample size based on the $N_{edge}$ value. Heterozygosity by loci was measured across 30 microsatellites by sampling 5 individuals per deme in 'scenario 1' from Fig. S1 ($d_{tot}$ = 382 demes) with $K$: 100, $m$: 0.04 and $\mu$: 5 x $10^{-4}$. Data points refer to one single simulation replicate. Dashed blue and solid black line indicate, respectively, theoretical results based on the estimated deme $N_{edge}$ and average value across 30 simulations. Solid thin line is the GAM model obtained from one simulation replicate. The insert bar plots quantify the percentage of variance explained by the 'edge-corrected population size' under the Wright-Malécot approximation (light blue) and GAM model (red). White diamonds indicate the theoretical $H_o$ using the Wright-Malécot approximation and ignoring edge-effect.

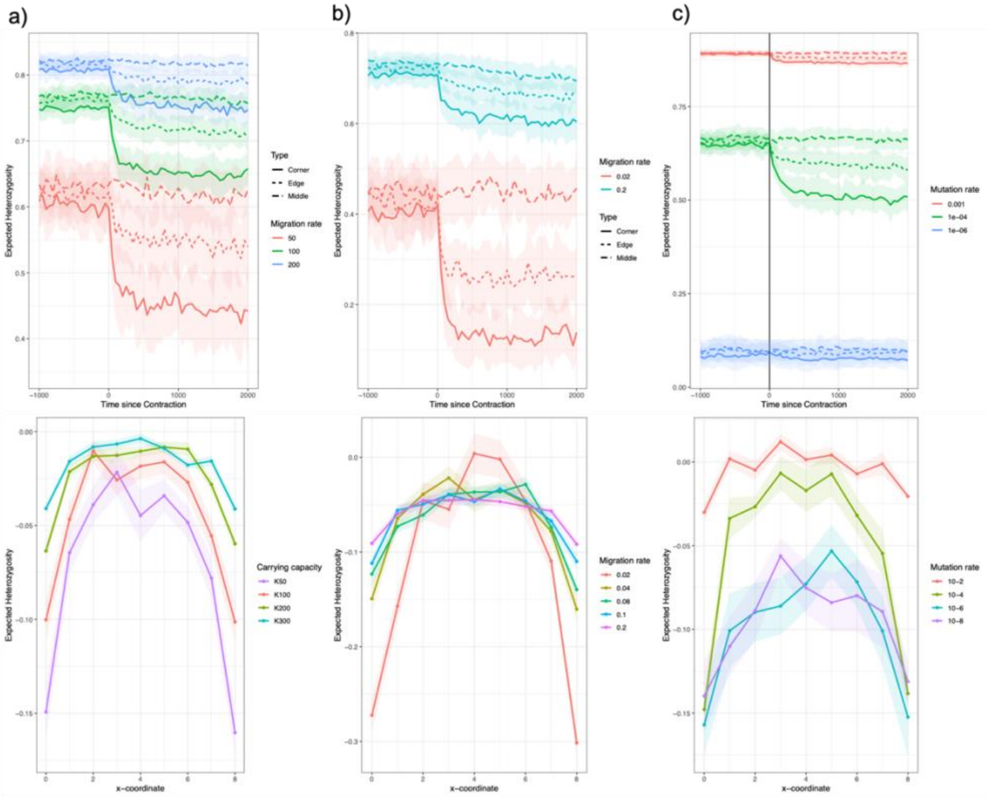


**Figure S12. Dynamic of deme $H_0$ in a 13 x 13 stepping-stone model undergoing habitat contraction.** Impact of **a)** carrying capacity (fixed parameters, $m$: 0.04; $\mu$: 5 x $10^{-4}$); **b)** migration rate (fixed parameters, $K$: 100; $\mu$: 5 x $10^{-4}$); and **c)** mutation rate (fixed parameters, $K$: 100; $m$: 0.04) on temporal changes in deme $H_0$. The first-row shows $H_0(t)$ for centre, edge and corner demes. The second row represents the difference in deme $H_0$ before and after contraction for demes sampled along the diagonal. Across all parameter's combinations, we sampled 14 individuals per deme, 30 microsatellite per individual. Dots and shade indicate, respectively, the mean and standard deviation of deme $H_0$ based on 30 simulation replicates.

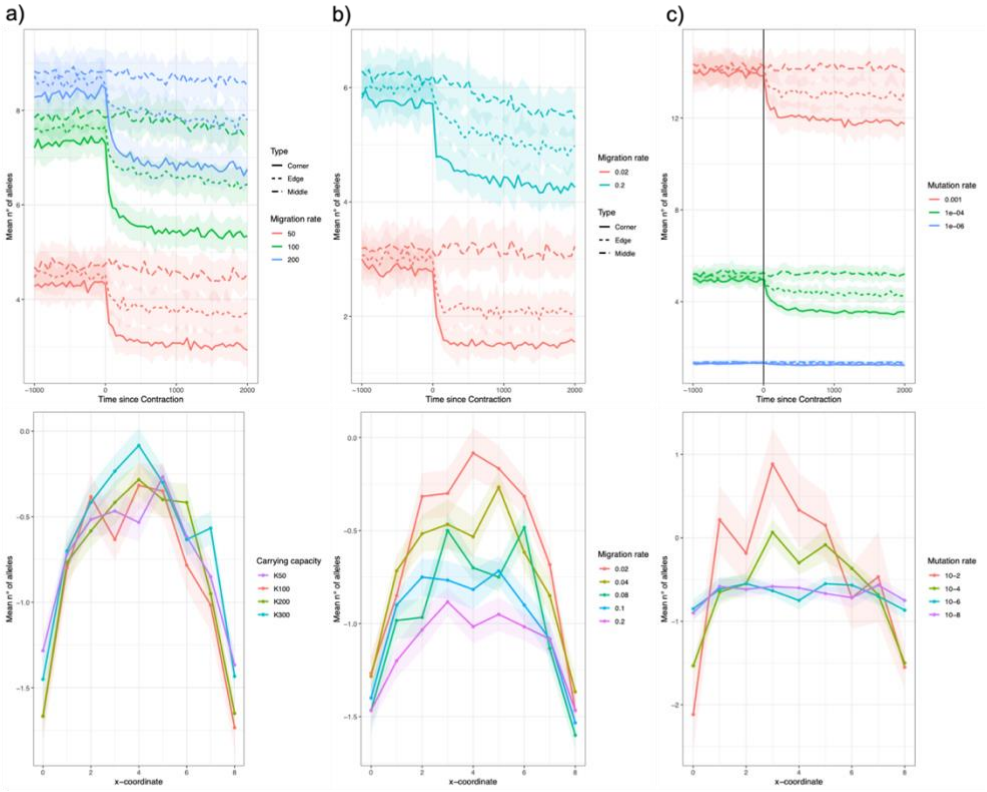


**Figure S13. Dynamic of deme $H_0$ in a 13 x 13 stepping-stone model undergoing habitat contraction.** Impact of **a)** carrying capacity (fixed parameters, $m$: 0.04; $\mu$: 5 x $10^{-4}$); **b)** migration rate (fixed parameters, $K$: 100; $\mu$: 5 x $10^{-4}$); and **c)** mutation rate (fixed parameters, $K$: 100; $m$: 0.04) on temporal changes in deme $H_0$. The first-row shows $H_0(t)$ for centre, edge and corner demes. The second row represents the difference in deme $H_0$ before and after contraction for demes sampled along the diagonal. Across all parameter's combinations, we sampled 14 individuals per deme, 30 microsatellite per individual. Dots and shade indicate, respectively, the mean and standard deviation of deme $H_0$ based on 30 simulation replicates.

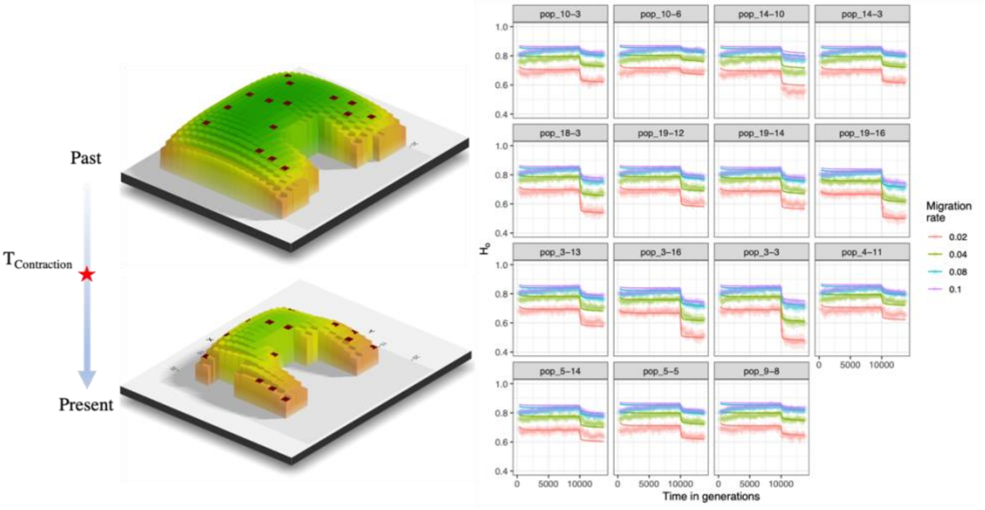


**Figure S14. Non-equilibrium deme $H_0$ in habitat patch (scenario 1) undergoing instantaneous habitat contraction.** Left column shows the simulated scenario. Colours and deme bars are proportional to the amount of edge-effect at which each deme is subjected to. Brown squares indicate sampled demes. Right column presents the comparison between simulation and analytical results for different values of migration rate, across demes subjected to various level of edge-effect. The solid lines represent theoretical predictions based on numerical integration of (*Eq. 8d*). Simulation results are represented by coloured dots and refer to the average deme $H_0$ across 30 simulation replicates, computed by sampling 14 individuals per deme at 30 microsatellite loci (fixed parameters, $K$: 100; $\mu$: 5 x $10^{-4}$). The results indicate good agreement between theoretical predictions and simulation results.

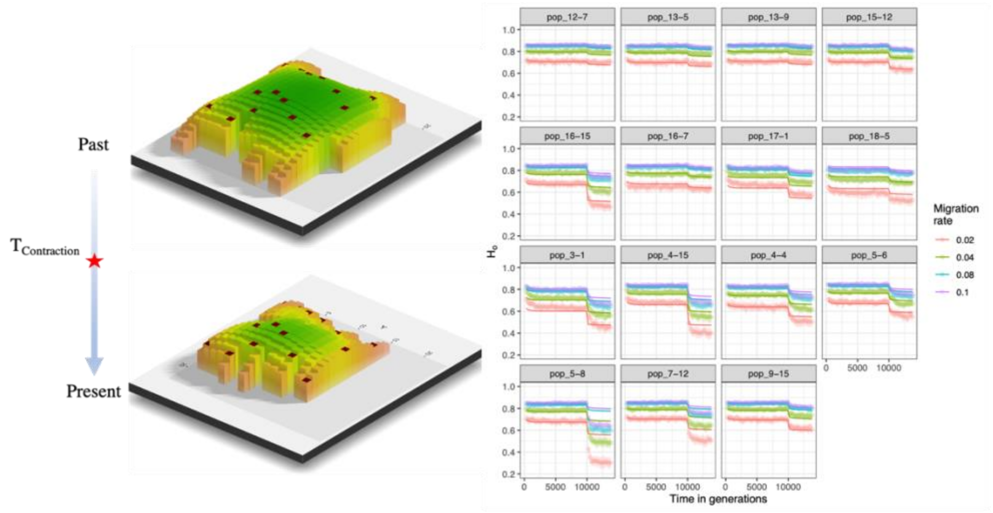


**Figure S15. Non-equilibrium deme $H_0$ in habitat patch (scenario 2) undergoing instantaneous habitat contraction.** Left column shows the simulated scenario. Colours and deme bars are proportional to the amount of edge-effect at which each deme is subjected to. Brown squares indicate sampled demes. Right column presents the comparison between simulation and analytical results for different values of migration rate, across demes subjected to various level of edge-effect. The solid lines represent theoretical predictions based on numerical integration of (*Eq. 8d*). Simulation results are represented by coloured dots and refer to the average deme $H_0$ across 30 simulation replicates, computed by sampling 14 individuals per deme at 30 microsatellite loci (fixed parameters, $K$: 100; $\mu$: 5 x $10^{-4}$). The results indicate good agreement between theoretical predictions and simulation results, except in few instancew (i.e., pop_5-8).

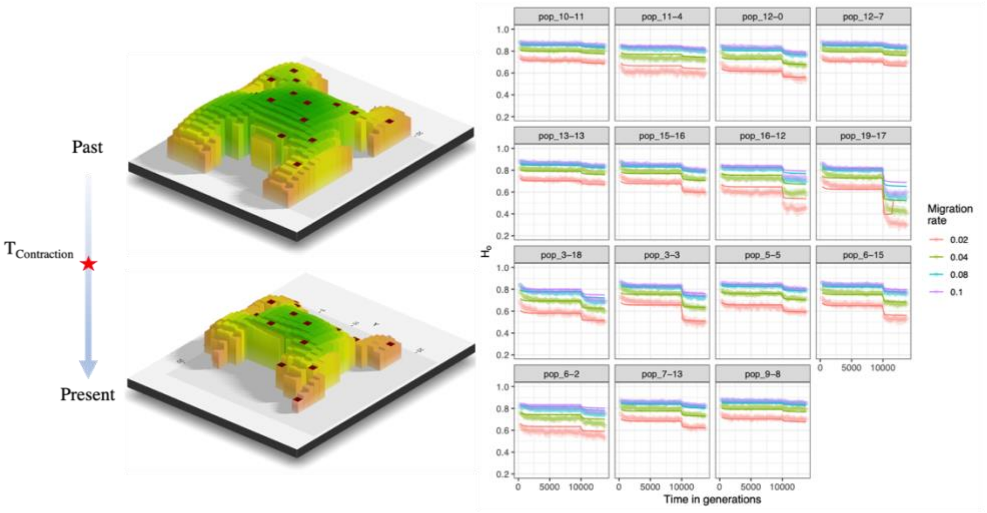


**Figure S16. Non-equilibrium deme $H_0$ in habitat patch (scenario 3) undergoing instantaneous habitat contraction.** Left column shows the simulated scenario. Colours and deme bars are proportional to the amount of edge-effect at which each deme is subjected to. Brown squares indicate sampled demes. Right column presents the comparison between simulation and analytical results for different values of migration rate, across demes subjected to various level of edge-effect. The solid lines represent theoretical predictions based on numerical integration of (*Eq. 8d*). Simulation results are represented by coloured dots and refer to the average deme $H_0$ across 30 simulation replicates, computed by sampling 14 individuals per deme at 30 microsatellite loci (fixed parameters, $K$: 100; $\mu$: 5 x $10^{-4}$). The results indicate good agreement between theoretical predictions and simulation results.

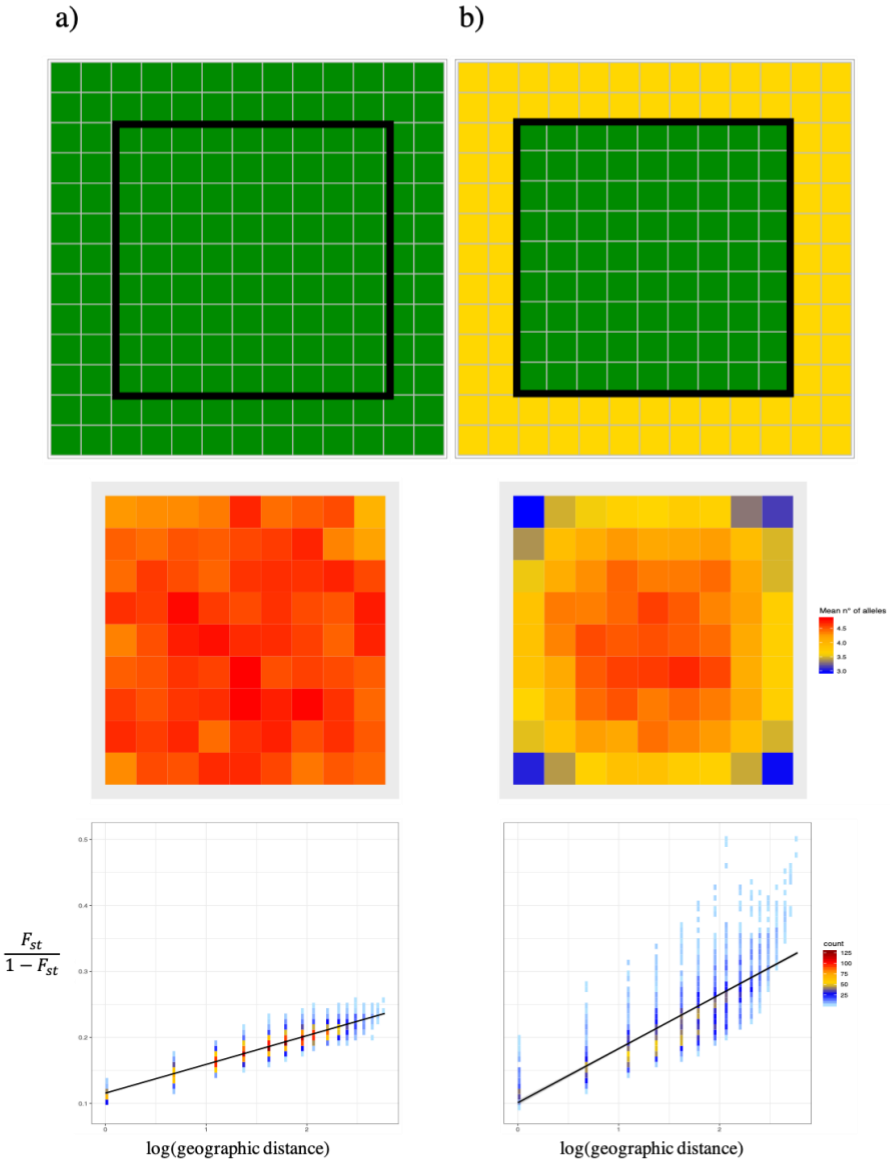


**Figure S17. The influence of edge-effect on genetic diversity and differentiation.** a) Before contraction; b) After contraction. First row: simulated scenario (13 x 13 demes) with $K = 100$, $m = 0.04$ and $\mu = 10^{-5}$, where dark green or yellow cells refer to habitat or no habitat. Dark line delimits the sampled demes. Second row: distribution of deme genetic diversity (MNA) across sampled demes. Third row: isolation-by-distance (IBD) pattern across all pairwise comparisons. The results show higher IBD slope after habitat contraction due to increase in edge effect. Five diploid individuals were sampled from each cell and simulated 30 microsatellite loci.

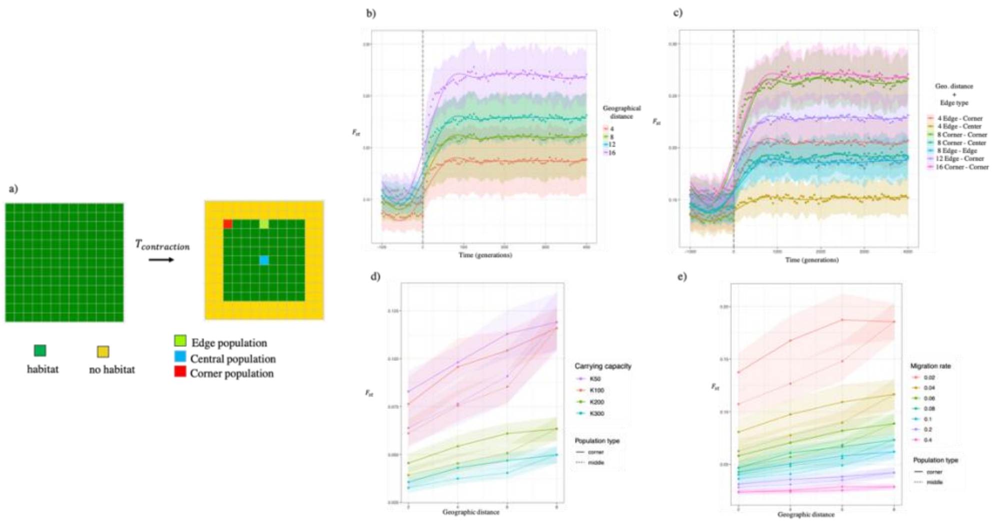


**Figure S18. The influence of edge-effect on isolation-by-distance.** a) simulated scenario (13 x 13 demes), where dark green or yellow cells refer to habitat or no habitat, while red, blue and light green cells correspond to corner, edge and central demes, respectively; b) Temporal dynamic of pairwise $F_{st}$ for deme pairs at different geographical distances (4, 6, 12, 16), dashed line represents the time of contraction; c) Temporal dynamic of pairwise $F_{st}$ for deme pairs at different geographical distances (4, 6, 12, 16) and including different edge types (Center, Edge, Corner), dashed line represents the time of contraction; d) the effect of 'Population type' on isolation-by-distance pattern, for several values of carrying capacity. Solid or dashed lines refer to pairwise comparisons including middle (=center) or corner demes, respectively. Panel b) and c) show that 'Edge type' can influence $F_{st}$as much as geographical distance, as shown for comparisons at distance 8. Panel d) and e) show that $F_{st}$comparisons including corner demes are higher than center ones.
